# Spatial Transcriptomics to define transcriptional patterns of zonation and structural components in the liver

**DOI:** 10.1101/2021.01.11.426100

**Authors:** Franziska Hildebrandt, Alma Andersson, Sami Saarenpää, Ludvig Larsson, Noémi Van Hul, Sachie Kanatani, Jan Masek, Ewa Ellis, Antonio Barragan, Annelie Mollbrink, Emma R. Andersson, Joakim Lundeberg, Johan Ankarklev

## Abstract

Reconstruction of heterogeneity through single-cell transcriptional profiling has greatly advanced our understanding of the spatial liver transcriptome in recent years. However, global transcriptional differences across lobular units remain elusive in physical space. Here, we implement Spatial Transcriptomics to perform transcriptomic analysis across sectioned liver tissue. We confirm that the heterogeneity in this complex tissue is predominantly determined by lobular zonation. By introducing novel computational approaches, we enable transcriptional gradient measurements between tissue structures, including several lobules in a variety of orientations. Further, our data suggests the presence of previously transcriptionally uncharacterized structures within liver tissue, contributing to the overall spatial heterogeneity of the organ. This study demonstrates how comprehensive spatial transcriptomic technologies can be used to delineate extensive spatial gene expression patterns in the liver, indicating its future impact for studies of liver function, development and regeneration as well as its potential in pre-clinical and clinical pathology.

## INTRODUCTION

The mammalian liver is a pivotal organ for essential metabolic homeostasis and detoxification. It has been ascribed a central role for the generation, exchange and degradation of essential biomolecules such as ammonium, fatty acids, amino acids and glucose as well as the conversion and eradication of various xenobiotic compounds and toxins^1^.

Depending on the species, the liver is divided into a specific number of lobes. In mice, the liver can be divided into four lobes: medial, left (largest), right (bisected) and caudate^2^. The mature liver architecture is arranged in repetitive units, termed liver lobules. In brief, the lobule, classically represented as a hexagon, has a portal vein (PV) at each junction to the neighboring lobules, through which blood rich in nutrients from the intestine enters into the liver. Eventually, the nutrient- and oxygen-exhausted blood is eventually drained in the central vein (CV) ^3–5^.

By area, the majority of liver resident cells (80%) are parenchymal cells, i.e. hepatocytes. The remaining 20% of the tissue consists of liver non-parenchymal cells (NPCs) including; liver endothelial cells (LECs), liver resident macrophages (Kupffer cells) and other immune cells, hepatic stellate cells (HSCs) and other stromal cells, biliary epithelial cells (cholangiocytes) and smooth muscle cells, which together make up the heterogeneous functional lobular liver environment ^6^. Hepatocytes execute distinct functions along the lobular axis based on their proximity to the CV or the PV ^7–10^. In mice, this spatial division in metabolic functions, known as zonation, is primarily based on the differential expression profiles of hepatocytes along the lobular axis and is classically divided into three zones, with zone 1 at portal veins, intermediate zone 2 and zone 3 at central veins ^10^. Recent findings from single-cell spatial reconstruction approaches suggest that smaller and less abundant NPCs also follow distinct spatial expression profiles based on their position along the lobular axis ^11,12^. These reconstruction approaches (1) provide an intricate image of the metabolic division of labor within the microenvironment of the liver lobule, (2) identify defining factors of zonation based on DGE along the lobular axis ^12–15^ and (3) represent a fundamental resource for the extensively studied concept of liver zonation ^6^. However, all previous studies either performed laser capture microdissection (LCM)^15^ or perfusion techniques ^12,14^, ultimately requiring tissue dissociation prior to sequencing, which is known to alter the physiological transcriptional landscape ^16–18^.

Further, previous studies focused on identifying factors underlying zonation exclusively in the microenvironment of the liver lobule. Investigation of individual liver sections shows that the theoretical organization of the repetitive liver lobules is challenging, due to the 3-dimensional organization and the overall complexity of the complete organ. Lobules across the tissue are organized in a highly irregular manner and differ greatly in size and axial orientation within the tissue. In addition, lobules are situated in varying proximities to the main sources of blood supply, namely the hepatic artery and the portal vein.

An additional layer of complexity studying liver tissues is created by their organization into several lobes ^19,20^. The reason for this partitioning is not yet fully understood, however, certain functional differences have been suggested ^21,22^. Gene expression profiles may also vary between regions, defined by their distance to other lobes. Therefore, DGE patterns among liver cells beyond their organization in individual lobules and in the extended tissue context are poorly studied and are vital for our full understanding of liver function in homeostasis and disease.

Spatial Transcriptomics (ST) enables high resolution assessment of spatial gene expression across tissue sections, overcoming the limitations associated with tissue dissociation ^16–18^. Hence, the generation of spatial transcriptomics data from liver sections in their bona fide tissue context, together with pre-existing knowledge of liver zonation enables spatial annotation of structures in the liver microenvironment (lobule) and liver macroenvironment (tissue section).

Moreover, performing ST across liver tissue sections has the capacity to reveal novel structures, which may be lost when using protocols that do not allow analysis in a spatial context -structures that may play crucial roles for the overall architecture of the liver.

Here, we perform ST on mouse liver tissue sections, assessing spatial factors contributing to spatial liver heterogeneity at the transcriptional level. By designing and implementing a variety of computational methods, this study aims to resolve the spatial relationships of vascular components involved in liver zonation and explore novel, previously uncharacterized structures based on their transcriptional profile and in their original tissue context. Our results support the concept that zonation represents the most prominent factor contributing to spatial heterogeneity. Computationally tracing the expression levels of genetic markers linked to zonation along the lobular axis allows us to study zonation gradients in physical space, and to infer the identity of vascular structures based on their expression profile. We anticipate that our results, combined with previous findings of different cell types constituting the overall transcriptional landscape of liver tissue, can enhance our current understanding of liver tissue organization.

## RESULTS

### Unsupervised clustering defines spatial distribution of expression across liver tissues

We used a total of 8 sections of wild type adult mouse livers from the caudate or right liver lobe for histological staining, library preparation and sequencing. After mapping, filtering, annotation and normalization we obtained expression data consisting of 19,017 genes across 4,863 individual capture locations (spots) on the ST array (summarized over all sections) and subjected the data to downstream computational analysis (see methods). Spots under the tissue section were considered for analysis and visualization (Figure 1a). Each spot is covered by a small mixture of liver cells. From the hematoxylin-stained nuclei, we estimate that each spot contains between 5-10 hepatocytes and up to 30 cells in total per spot. Subsequently, we embedded the data into low-dimensional space via non-negative matrix factorization and clustered it in an unsupervised manner using a graph-based approach, which identified 6 clusters (Figure 1b, top panel, see methods for details). To put the clusters into context and assess their spatial organization, spots were projected on the brightfield image of Hematoxylin- and Eosin (H&E) stained tissue.

**Figure 1.**
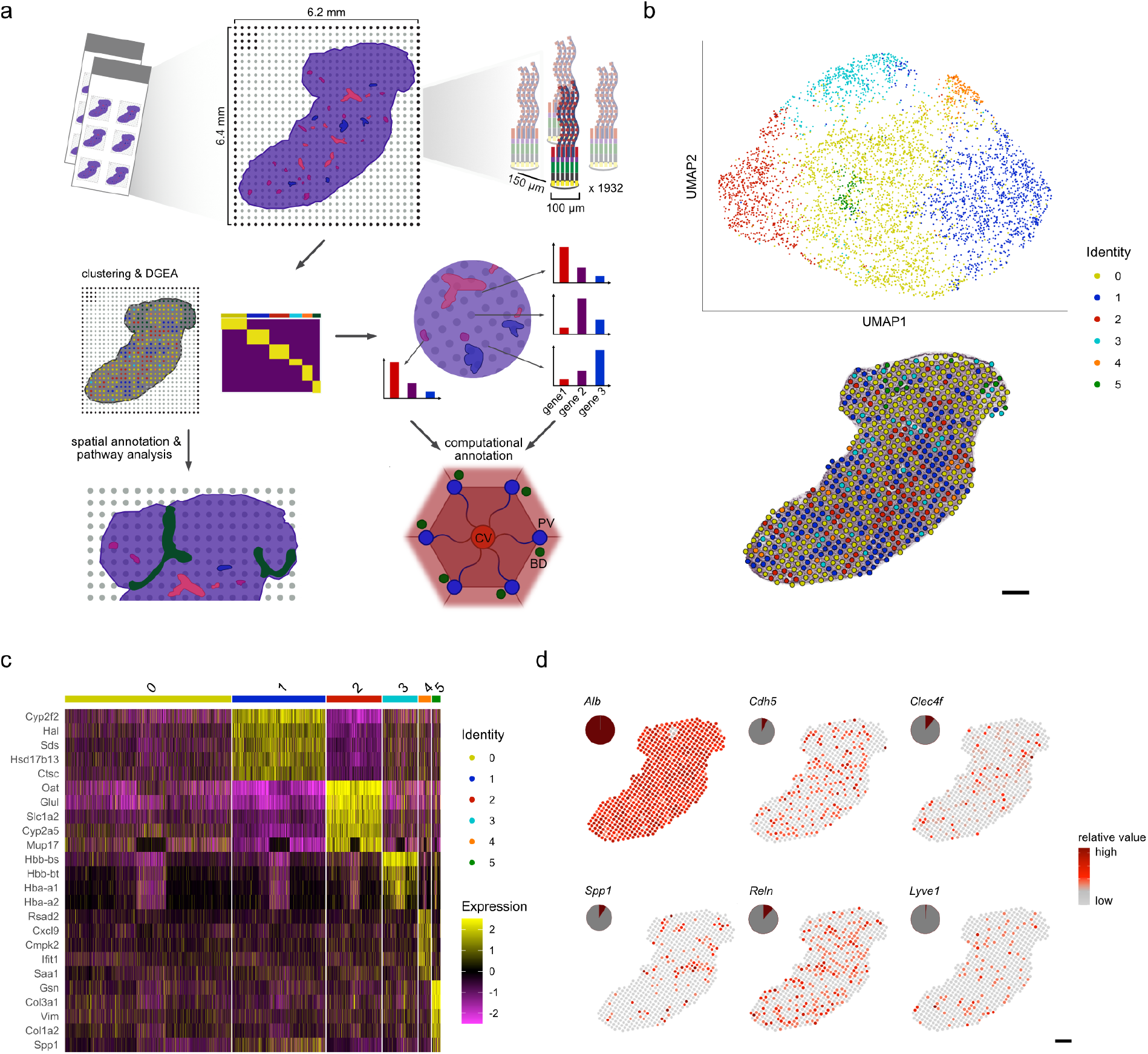
Study overview of spatial transcriptomics on murine liver. **a)** Spatial transcriptomics was performed on a total of 8 murine liver tissue sections. The tissue sections were placed in one of six, 6.2 x 6.4 mm frames on the glass slide ST array. Each frame contains 1932 spots, with >200M uniquely barcoded, mRNA capture probes. The distance between centers of each neighboring spot is 150 µm. Initially, each tissue section was fixed, stained with hematoxylin and eosin (H&E), followed by imaging. Tissue sections were permeabilized, followed by mRNA capture, tissue removal and sequencing. Thereafter, the count data was subjected to cluster- and differential gene expression analysis (DGEA). The results of the clustering and DGEA were further analyzed and spatially annotated at the global tissue context and down to the lobular level. For new spatial annotations, pathway analysis was performed. Liver lobules are classically described by a central vein (CV, red) surrounded by 6 portal nodes (PV, blue) with neighboring bile-ducts (BD, green). For lobular spatial annotations, clusters have been computationally annotated by comparing expression levels in a set of genetic markers linked to metabolic zonation along the lobular axis. **b)** Canonical correlation analysis (CCA) was performed to integrate data of eight liver tissue sections, the data was subsequently normalized and subjected to graph-based clustering in which 6 clusters were identified (see methods). The integrated data was embedded in UMAP space (top) and depicted as an overlay of the spot cluster annotation across the tissue (bottom) (scale bar indicates 500 µm). **c)** Heatmap depicting expression values of the five most variable genes for each cluster after subjecting the six clusters to DGEA, with the exception of cluster 3, which resulted in only four significantly differentially expressed genes. **d)** Visualization of spatial distribution of reported expression markers of Hepatocytes (*Alb*), liver endothelial cells (*Cdh5*), Kupffer cells (*Clec4f*), Cholangiocytes (*Spp1*), hepatic stellate cells (*Reln*) and lymphatic liver endothelial cells (*Lyve1*) by spots under the tissue. Pie-charts indicate the respective proportion of cell type markers present in spots under the tissue (scale bar indicates 500 µm).

The projection showed a clear spatial segregation between spots belonging to certain clusters. At a first glance, cluster 5 was localized to an exclusive region of the tissue section, while spots belonging to cluster 1 and cluster 2 visually seemed to align with the vascular structures in the liver tissue (Figure 1b, lower panel, Supplementary figure 1). To further describe the identified clusters, we performed DGE analysis between them. In fact, differentially expressed genes (DEG) in cluster 1 support periportal gene expression from previous studies while genes previously associated with pericentral gene expression are enriched in cluster 2, proposing that cluster 1 and cluster 2 denote regions around portal and central veins, respectively ^10,11,15,16^. Cluster 3 shows enrichment for genes associated with hemoglobin, whereas cluster 4 shows enriched expression of genes involved in immune-related processes ^23,24^. Cluster 5 displays enrichment for mesenchymal genes ^25–27^ (Figure 1c, Supplementary table 1).

Spots of cluster 3, cluster 4 and cluster 5 are mainly surrounded by spots of different clusters, while cluster 0, 1 and 2 form more cohesive groups of spots. Interestingly, spots of cluster 0, 3 and 4 seem to adjoin spots of cluster 0, 1 and 2 in descending order, implicating transcriptional profiles of most clusters are commonly surrounded by periportal rather than to pericentral areas (Supplementary figure 2).The scattered spatial distribution of cluster 3 across sections can most likely be explained by the fact that the tissue was not perfused prior to freezing and sectioning, allowing us to detect blood cell populations throughout the liver. To approximate the replicability and the sensitivity of the method to detect the expected transcriptome of liver cells per spot, we examined the expression of genes, reported to be markers for common cell types in liver across spots under the tissue.

**Figure 2.**
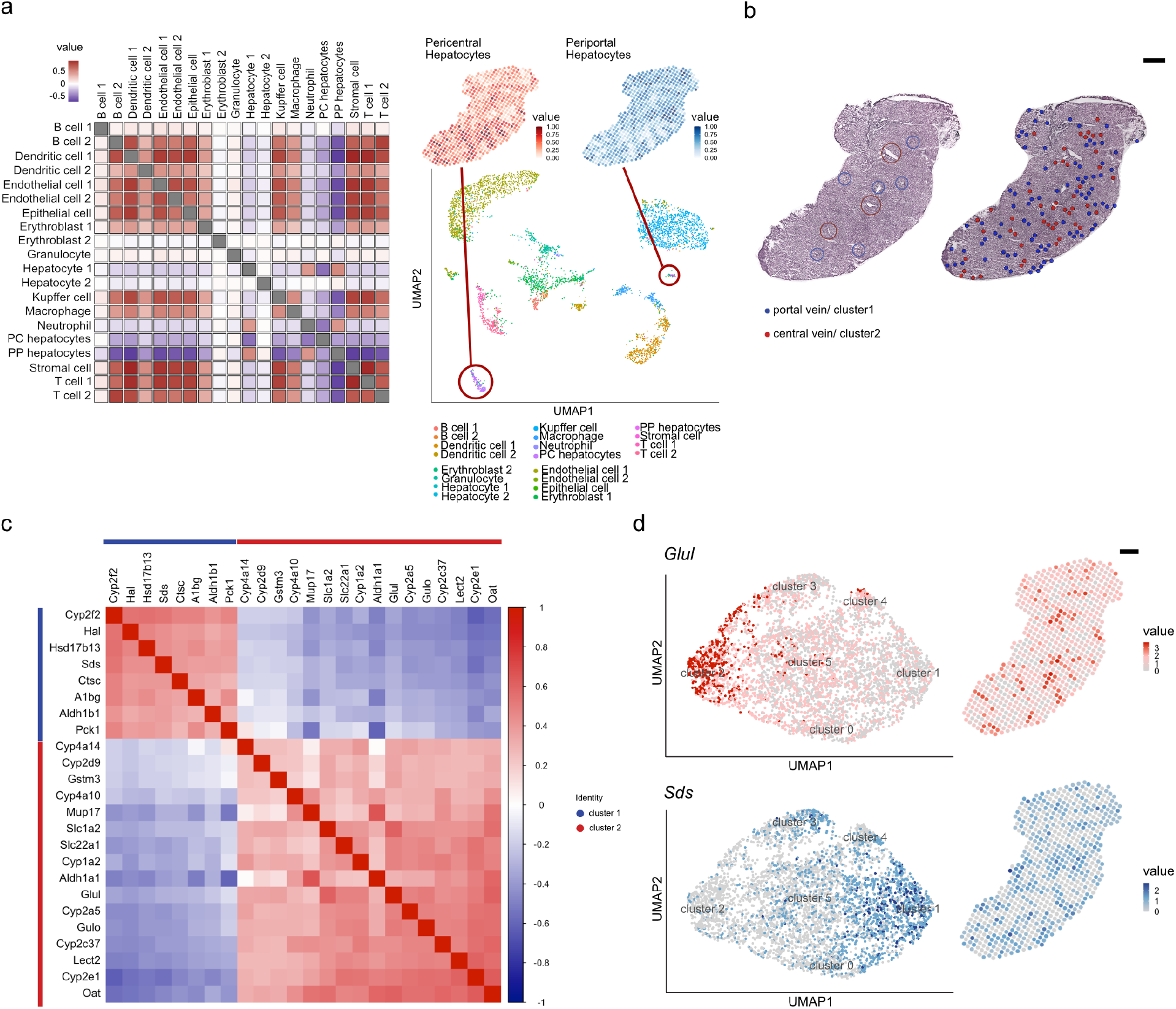
Clustering, spatial annotation and computational validation using established scRNA-seq data. **a)** Visualization of cell type co-localization by Pearson correlations (left). Positive correlation values indicate spatial co-localization of cell types while negative values represent spatial segregation. UMAP embedding of single-cell data of the Mouse Cell Atlas (MCA)^41^ grouped by annotated cell types (bottom right). Numeration behind the cell types represent annotation of MCA data (B cell-1 = Fcmr high, −2 = Jchain high, Dendritic cell-1 = Cst3 high, −2 = Siglech high, Epithelial cell-1 = Spp1 high, −2 = /, Eryhroblast-1 = Hbb-bs high, −2 = Hbb-bt high, Hepatocyte-1 = Fabp1 high, −2 = mt-Nd4 high, T cell-1 = Gzma high, −2: Trbcs2 high). Encircled clusters in the plot refer to pericentral or periportal hepatocytes of MCA data. Quantile scales of cell-proportions annotated as pericentral and periportal hepatocytes (see methods) are mapped on spatial transcriptomics spot data (top right). **b)** Visualization of spots representing gene expression profiles of cluster 1 (portal vein, blue) and cluster 2 (central vein, red) on H&E stained tissue (right), compared with visual histology annotations of central-(red circles) and portal-(blue circles) veins (left) (scale bar indicates 500 µm). **c)** Pearson correlations of genes expressed in cluster 1 and 2 ordered by their first principal component (see methods). Genes with high expression in the pericentral cluster (cluster 2) show negative correlation with genes highly expressed in the periportal cluster (cluster 1) and vice versa. Genes present within cluster 1 or cluster 2 exhibit positive correlation with genes in the same cluster. **d)** Projection of selected markers for central venous expression (*Glul*, top) and periportal expression (*Sds*, bottom) in UMAP space and spots under the tissue (scale bar indicates 500µm).

In alignment with the histological evaluation of the tissue, expression of the hepatocyte marker *Alb* (expression value > 1) in all 4863 spots (100%) indicates all spots contain hepatocytes. For LECs, 1972 spots showed expression of *Cdh5* ^28,29^ (∼8%). Lymphatic liver endothelial cell marker *Lyve1* ^30–32^ showed expression in a small fraction of only 80 spots (0.02%). Kupffer cell marker *Clecf4* ^33–35^ showed expression in 526 spots (∼11%) while stellate cell marker *Reln* ^36^ was expressed in 568 spots (12%). *Spp1* is a marker for Cholangiocytes ^37^, expected to only be present in bile ducts, next to portal veins and is expressed in 452 spots (∼9%) (Figure 1d). These results demonstrate that cells with larger volumes and higher abundance in the liver are represented as such in the data, whereas smaller and rarer cell-types are found more scattered across the tissue, as would be expected.

While characteristic marker gene expression is a common way to extrapolate the presence of certain cell types, we wanted to include a larger set of genes constituting the expression signature of a specific cell type and compare it to our spatial data. *Stereoscope*, developed by Andersson et al. ^38^ enables cell types from single cell RNA sequencing (scRNA-seq) data to be mapped spatially onto the tissue, by using a probabilistic negative binomial model. Using this method, we were able to spatially map 20 cell types annotated by the Mouse Cell Atlas (MCA) ^39^ on liver tissue sections (Supplementary figure 3.1 - 3.3). Notably, high proportion values are estimated for periportal as well as pericentral hepatocytes in the MCA (Supplementary figure 3.1 − 3.3). Pearson correlation scores between cell type proportions across the spots show positive correlation, to be interpreted as spatial co-localization of non-parenchymal cells like LECs, epithelial cells and most immune-cells, as well as stromal cells (Figure 2a). For cell types related to immunity, only neutrophils display slight positive correlation to periportal hepatocytes and *Fabp1-*enriched hepatocytes, simultaneously exhibiting negative correlation to pericentral hepatocytes, indicating spatial segregation. Interestingly, periportal and pericentral hepatocytes not only exhibit negative correlation between each other, but also to most other cell types (Figure 2a). A large portion of spots is assigned to cluster 1 and cluster 2, and 100% of the spots contain hepatocyte markers, showing that - spatially - the liver is predominantly constituted by zonated hepatocytes, while these cells only represent a very small fraction of the MCA data. This discrepancy illustrates the power of complementing single cell trancriptome data with spatial gene expression data to thoroughly delineate liver architecture and the transcriptional landscape of liver tissue, while simultaneously demonstrating the limits of scRNA-seq data integration. Importantly, correlations between periportal and pericentral hepatocytes and portal and central clusters show similar trends as observed for spatial data, advocating for a reliable integration of cell type annotations from scRNA-seq data and our ST data (Figure 2a, Supplementary figure 4, Supplementary table 3.2).

**Figure 3.**
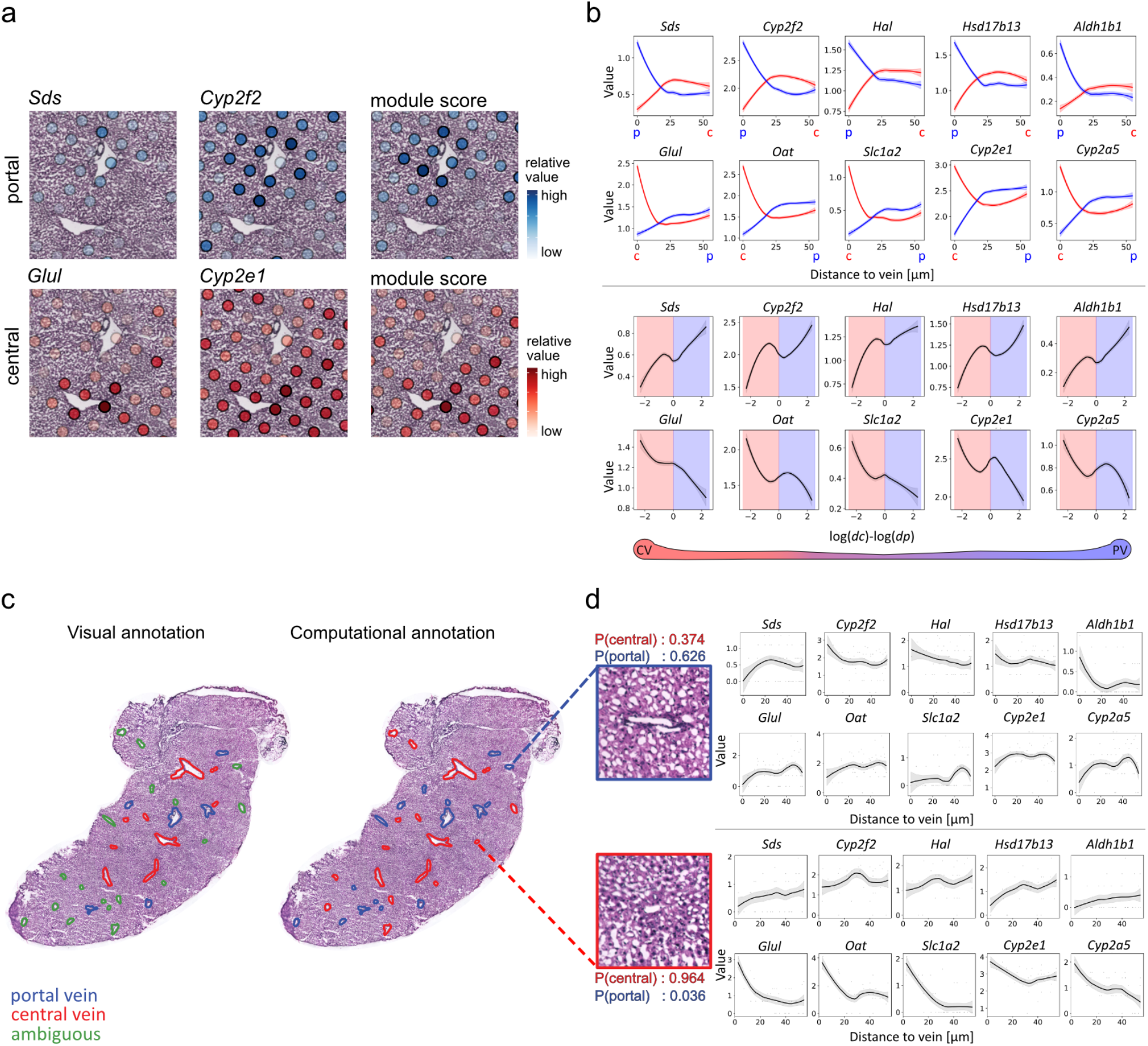
Expression gradient along the lobular axis and computational annotation of liver vein types. **a)** Enlarged view of a superimposed visualization of *Sds, Cyp2f2* expression in the portal vein module, consisting of selected DEGs of cluster one (supplementary table 1), all with high values around the histological annotation of a portal vein (top). Expression of *Glul, Cyp2e1* as representative marker-genes of the central vein module expression (supplementary table 1), consisting of DEGs of cluster 2 with high values around the histological annotation of a central vein (bottom). **b)** Visualization of the average expression by distance to vein-type measured within 50 µm from the vein. The top row shows expression by distance of portal markers *Sds, Cyp2f2, Hal, Hsd17b13* and *Aldh1b1* to portal veins in blue and central veins in red, while the bottom row shows distances of central vein markers *Glul, Oat, Slc1a2, Cyp2e1* and *Cyp2a5* to portal veins in blue and central veins in red (top panel). Visualization of relative proximity of portal vein markers in the top row and central vein markers in the bottom row to both vein types. The gene expression as a function of the logged relative distances (see methods for details), negative values on the x-axis indicate points with closer proximity to central veins in red compared to portal veins in blue and vice versa for positive values, as indicated by the schematic (below graphs). **c)** Visual histological annotations (left) of central (red) and portal (blue) veins, including ambiguous visual annotations (green), compared with computational prediction, using the 10 marker genes from 3b (right). The classification of vein types is based on a weighted (by distance) average expression of the genes expression profiles in the neighborhood of each vein. In addition, the spatial expression data of spots neighboring uncertain morphological vascular annotations (green) can be used to deduce periportal or pericentral vein-types in the cases where visual annotations are ambiguous. **d)** Expression by distance of portal - (top panel) and central - (bottom panel) markers. Probabilities for each class (central and portal) can be extracted from the logistic regression model, here given as P(central) or P(portal) (scale bar indicates 500µm).

**Figure 4.**
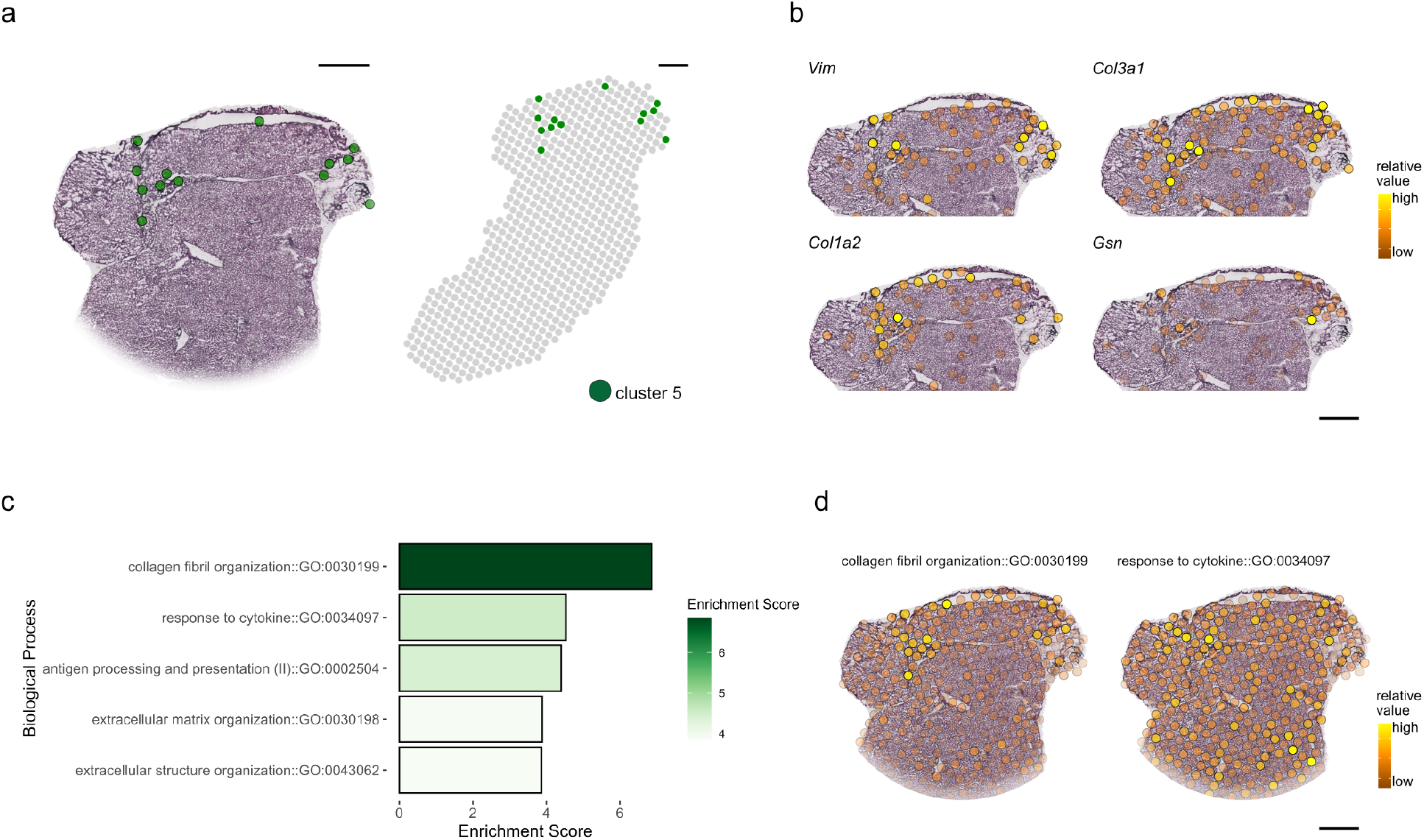
Identification of liver tissue regions with unique transcriptional patterns. a) Projection of spots including transcriptional patterns of cluster 5 in the UMAP (from Fig 1b), on a selected part of a histological section of the caudate lobe (left) and spot location in the entire tissue section (right). b) Visualization of *Vim, Col3a1, Col1a2* and *Gsn* expression in spots of the same tissue section as in 4a. c) Gene-ontology (GO) enrichment for markers present in cluster 5. The Enrichment is given as the negative log10 algorithm of the adjusted p-value” (g:SCS correction, see methods) of the differentially expressed marker genes in cluster 5. d) Module scores of genes (see methods) in cluster 5 belonging to the two biological processes with the highest enrichment scores: “collagen fibril organization” and “response to cytokine” are visualized on spots across the tissue.

### Heterogeneous spatial gene expression linked to pericentral and periportal zonation

Spatial mapping of common marker genes of periportal or pericentral zonation as well as periportal and pericentral hepatocytes from single-cell integration imply co-localization of cluster 1 and cluster 2 with portal and central veins respectively. To support this observation, venous structures in our sections were annotated as a portal vein, central vein or vein of unknown type (ambiguous) based on the presence of bile ducts and portal vein mesenchyme (PV) or lack thereof (CV). Comparison of the histological annotations and the corresponding clusters sustain the observation that cluster 1 denotes periportal areas and cluster 2 signifies pericentral areas in the tissue sections (Figure 2b).

Pearson correlations between genes enriched in periportal cluster 1 (PPC) and genes enriched in pericentral cluster 2 (PCC) show a negative trend, interpreted as spatial segregation (Figure 2c, Supplementary table 2). PCC genes exhibit positive correlations to all other marker genes present in the PCC, and PPC marker genes show positive correlations to other PPC markers, interpreted as spatial correlation (Figure 2c). None or lower correlations can be observed between PPC or PCC marker genes and the remaining 4 clusters (cluster 0, cluster 3, cluster 4 and cluster 5) (Supplementary Figure 5, Supplementary table 2). The observed heterogeneity of spatial gene expression across liver tissue sections based on portal or central vein-identity is further supported by spatial autocorrelations of marker genes, where values higher than 0.2 were considered to exhibit autocorrelation. Higher values indicate stronger positive correlation of gene expression and tissue localization, while lower values (<0.2) suggest independence of gene expression and space (Supplementary Figure 6, supplementary table 3).

Visualization of spots with expression of representative pericentral (*Glul*) and periportal (*Sds*) markers in the UMAP embedding further demonstrate highest expression values of *Glul* or *Sds* in the pericentral or periportal cluster, respectively. Thereafter, the expression values decrease exhibiting the lowest values for spots in the cluster representing the opposite vein-type. When displaying expression of *Glul* and *Sds* on spots in their spatial context, these genes show highest expression in areas annotated as central or portal veins. In addition, no expression of *Sds* can be found in areas of high *Glul* expression and vice versa, indicating expression of genes present in the pericentral cluster 1 and periportal cluster 2 are spatially distinct and negatively correlated with each other (Figure 2d).

### Transcriptional profiling of pericentral and periportal marker genes across tissue space enable computational annotation of liver veins

To further investigate zonation in physical space, we first superimposed the spots under tissue showing expression for two representative markers of central veins (*Glul, Cyp2e1*) and portal veins (*Sds, Cyp2f2*), on histologically annotated veins (Figure 3a). *Glul* is responsible for the expression of glutamine synthase, the main enzyme for glutamine synthesis ^13^, while serine dehydratase (*Sds*) is a key factor for gluconeogenesis ^40^. *Cyp2e1* and *Cypf2* both belong to the cytochrome p450 family involved in xenobiotic metabolism ^41–43^. Pericentral expression of *Glul* is restricted to spots in very close proximity to an annotated central vein, while *Cyp2e1* is more evenly distributed across spots, with both genes not detectable around a nearby annotated portal vein. Similar observations are made for the expression of *Sds* and *Cyp2f2* around the portal vein. Including all marker genes of the PCC and the PPC and creating module scores of expressions of all DEG of the respective cluster in the spots under tissue, we visualize the common expression gradient along the lobular axis (Figure 3a, Supplementary figure 7).

Next, we wanted to assess whether gene expression was influenced by spatial proximity to the different vein types, as would be expected based on the study by Halpern et al., describing expression gradients over a total of 9 layers along the lobular axis ^14^. For this purpose, we generated what will be referred to as *expression by distance* plots; which portray the normalized gene expression as a function of the distance to respective vein type. To construct these plots, for each spot and gene, we pair the observed expression value with the distance from the spot’s center to the nearest vein border. Finally, to better capture the relationship between distance and expression, we smooth our observations with the *loess* method (see methods). Expression by distance plots were compiled for a select set of five periportal and pericentral marker genes with the highest positive logFC in the PPC and PCC (Supplementary table 1, Supplementary figure 14). Upon inspection of the plots, a clear dependency between distance and expression become apparent. Portal markers exhibit a gradual decline upon increased distance from a portal vein. For central veins, certain genes (e.g. *Glul, Slc1a2 and Oat*) exhibit a steep decrease in expression as the distance from a central vein increased, while others (e.g., *Cyp2e1* and *Cyp2a5*) display a more gradual decline (Figure 3b). These results are in agreement with the observed expression gradients in spatially reconstructed layers in Halpern et al ^14^ (Supplementary figure 8). Additionally, we aligned the mapped proportion-values of the annotated periportal- and pericentral hepatocytes in the MCA single-cell data along the lobular axis, and observed the same inverse relationship with the distance of their associated vein types as was observed for the marker genes. (Supplementary figure 9).

While the expression by distance plots disclose the influence of one vein type on gene expression in physical distance, we wanted to account for the eventual proximity to both vein types simultaneously. Hence, we developed expression by distance-ratio plots, where the logged ratio between the distance of both vein types is used. Using these proportion values of the physical distance of the expression by distance plots, we can account for the presence of both vein types, investigating their impact on the spatial expression landscape. We observed an almost linear relationship for portal genes along the lobular axis, exhibiting the lowest values closest to central veins and highest marker gene expression closest to portal veins. Meanwhile, expression of central markers exhibits a steep negative slope in very close proximity to central veins with a less steep decline towards portal vein vicinity (Figure 3b). The observed differences in expression along the lobular axis of different central and portal vein markers agree with early concepts of “dynamically” expressed genes along the lobular axis of the “gradient” type and “stable” gene expression of genes of the “compartment” type directly at the central or portal vein borders ^8,44,45^. Spatially stable expression of compartment type genes is exemplified by *Glul*, and glutamate transporter *Slc1a2*, important for glutamine transport at central veins ^13^. Expression of *Sds*, and the histidine ammonia lyase (*Hal*), involved in ammonium production are distinctive for stable gene expression at portal veins ^46^. The dynamic expression of gradient type genes is illustrated by *Cyp2e1* (pericentral) and *Cyp2f2* (periportal).

Given the strong association between the DEGs in the PPC and PCC, as well as the convincing demonstration of co-localization with histologically annotated central and portal veins, we aimed to explore the potential to annotate central and portal veins computationally, solely based on gene expression (Figure 3c). Computational annotation of veins as a complement to manual annotation is of relevance for multiple reasons. First, visual annotations sometimes prove to be difficult when only histological images of suboptimal quality or without immunohistological staining are available. Secondly, it is a labour intensive process that requires thorough histology training, which is not always available. Thus, a computational model not only provides the possibility to support visual vein annotations but also to predict the type of unannotated veins based on their surrounding gene expression profiles. The model constructed in this study (see methods) corresponds convincingly to visually annotated central and portal veins based on the expression profile of their respective neighborhood across all sections from different biological origin (caudate- and right liver lobe). Based on the confident attestation of overlapping visual- and computational vein annotation, we continued to computationally annotate veins with uncertain identity. Our results show estimated vein-assignment of 72 ambiguous veins to either central- or portal veins, inferred from the expression by distance of a subset of 5 central- or portal vein markers along the neighborhood of each vein (Figure 3d, Supplementary table 5). The deduction of vein-types based on the spatial expression profile of surrounding spots demonstrates the potential to use spatial gene expression data for a variety of annotation-based applications.

### Exploration of components contributing to spatial heterogeneity across liver tissues

Projecting the spot coordinates assigned to cluster 5 on the stained images demonstrates exclusive spatial organization in one or two distinct regions across the tissue (Figure 4a, Supplementary figure 10). Therefore, we asked how this cluster fits into the spatial liver organization based on its expression profile. Additionally, we wanted to assess whether the spatial organization of this cluster can give indications regarding the function of the underlying distinct morphological structure in this region of the tissue, which is characterized by morphologies resembling potential tissue partitioning.

DGEA identified *Gsn, Col1a2, Col1a3* and *Vim*, as highly upregulated marker genes of cluster 5 (Supplementary table 1). Spots outside cluster 5 identity, show no expression or low expression of these genes is observed (Figure 4b, Supplementary figure 11). Of the four genes, *Col1a2* and *Col3a1* indicate the highest expression upon visual inspection. In fact, pathway analysis of the cluster 5 marker genes demonstrates the strongest enrichment of genes belonging to the “collagen and fibril organization” gene ontology (Figure 4c, Supplementary table 6). Apart from *Col1a2* and *Col3a1* four more marker genes in cluster 5 belong to this biological process (*Dpt, Col1a1, Lum* and *Col14a1*). Collagen-fibrils have been reported to be the main component of the irregular connective tissue composing the Glisson’s capsule in several animals, including rodents ^47^, giving first indications for a structural function of cluster 5. In addition, processes contributing to structural formation and development, such as “extracellular matrix organization” and “extracellular structure organization” and pathways related to innate immunity, namely “response to cytokine”, “antigen processing and presentation of peptide or polysaccharide antigen via MHC class II” show enrichment within cluster 5 (Figure 4c, Supplementary table 6).

Expression scores of markers involved in “collagen and fibril organization” are highest in spots in spots of cluster 5 and in their direct proximity in the tissue and show low scores for the remaining tissue. In contrast, expression scores of marker genes involved in the response to cytokines (*H2-Eb1, Timp2, Timp3, H2-Aa, Cd74, H2Ab1, Spp1, Gsn, Col3a1, Vim*) are more evenly distributed across the tissue (Figure 4d, Supplementary figure 12). This result supports the higher significance of processes involved in structural formation and development of the tissue area at and around cluster 5.

Moreover, Pearson correlations between marker genes of all clusters demonstrate that most cluster 5 markers show no correlation between genes of cluster 1 or cluster 2 (Supplementary figure 5) and a majority of cluster 5 markers shows significant spatial autocorrelation. Taken together, correlation analysis of cluster 5 markers and histological morphology in the respective tissue area advocates for the spatial organization of cluster 5, independent of liver zonation (Supplementary figure 6, Supplementary table 3).

## DISCUSSION

Applying Spatial Transcriptomics on the mammalian liver represents a novel, compelling venue to explore its transcriptional and functional heterogeneity while also complementing previous data ^6,17^. Recent scRNA-seq studies including spatial integration by reconstruction provide high resolution information of single cell transcriptomes ^13,14,14,15^, but the spatial composition of these cells within the tissue is lost due to tissue dissociation, which additionally increases the risk of undesirable transcriptional changes ^17,18^ . In contrast, ST preserves the spatial information of gene expression in their true tissue context, thus categorically complementing single cell transcriptomic approaches. The emerging possibilities of combining spatial transcriptomic data with de-novo and existing single cell and other omics data of the same tissue offer unprecedented levels of insight into the biology of the tissue ^38,48^.

Here, we incorporated cell type information in the spatial data in two different ways. First, we assessed expression of characteristic marker genes within a wide range of expression levels. One example of a gene, expected to be sparsely present, is the lymphatic vessel endothelial hyaluronan receptor (*Lyve1*). In contrast to these rare cell types, liver tissue is reported to consist of up to 80% of hepatocytes, by area ^6^. Accordingly, *Alb* transcripts were detected at high frequency in all spots under the tissue. A recent study suggests predominant localisation of Kupffer cells in the periportal area of the liver lobule and Neutrophil recruitment upon bacterial infection ^49^. While, our data does not indicate significant enrichment of Kupffer cell markers in the periportal cluster of our data, colocalization of neutrophils and periportal hepatocytes, suggests periportal localization of neutrophils already in unperturbed conditions supporting implications of a proposed immune zonation ^49^. The liver is constantly exposed to toxic and microbial threats from the periportal blood, requiring an efficient balance between immune hyporesponsiveness and effective clearance of pathogens ^50^. Therefore, it will be of high interest to perform spatial transcriptomics to study the effect of infection and inflammation on the proposed immune zonation.

Secondly, deconvolution of mixed cell expression profiles using *stereoscope* shows higher proportion values for pericentral and periportal hepatocytes than all other cell types, potentially explaining the predominant spatial segregation between pericentral and periportal hepatocytes and most other cell types in our data. The observed discrepancies between hepatocyte numbers in our and the MCA data may result from the different technical limitations that scRNA-seql as well as spatial data generation face, emphasizing the current limits of scRNA-seq data integration. Transcripts from transcriptionally highly active or physically large cells might mask cell types with moderate to low transcriptional levels. Therefore, technical and computational advances to enhance resolution may benefit transcriptional profiling of rare cell types within a tissue. Nevertheless, comparisons to scRNA-seq data confirm general trends observed in our ST data, highlighting the importance of combining spatial transcriptomics with scRNA-seq data.

We annotated two clusters with anti-correlating spatial distributions and characteristic marker gene expression that align well with visually annotated portal or central veins in the H&E image as periportal (PPC) and pericentral (PCC) clusters. Overall, the spatial data generated in this study supports the hypothesis that the main source of spatial heterogeneity across liver tissue are transcriptional differences between zones along the lobular axis between portal and central veins ^13,12,14^.

Further, exemplary expression of “compartment” type zonation central markers *Glul* and *Slc1a2* and portal markers *Sds* and *Hal* illustrate that compartmentalization of gene expression for genes performing opposing tasks like glutamine and ammonium synthesis is necessary to prevent futile cycles ^51^ and illustrates one example of the relevance of biochemical zonation along the porto-central axis, which is thoroughly reviewed ^5,6,8,10,12–15,52,53^. Our data introduces distance thresholds for “compartment” and “dynamic” markers and traces expression gradients from outer vein borders and across physical space.

In addition, we investigate the relationships between marker gene expression of both portal and central veins simultaneously. Veins annotated as being central in the tissue depict a convincing inverse relationship between expression and distance to the portal vein type. Marker gene expression across all visually annotated veins in the tissue is insufficient to confirm the proposed schematic organization of the liver lobe of one central vein surrounded by six portal nodes. Nevertheless, the results implicate that the overall relationships of zonation markers between central and portal veins are comparable between porto-central axes across the tissue, independent of the schematic organization of lobules in physical space.

Based on the compelling evidence for robust expression profiles of central and portal veins across the tissue we were able to generate a computational model to predict the vein-type in cases where visual annotations were ambiguous, based on the expression profiles of neighboring spots. This computational model demonstrates the great potential of spatial transcriptomics to assist morphological annotations, providing probability values for the certainty of the computational annotation of morphological structures at their natural tissue location by transcriptional profiling. We anticipate that this method will provide a multitude of applications in future spatial transcriptomics studies, e.g. linked to pathology or infection.

Cluster 5 consists of a small number of spots with distinct spatial localization. As a consequence, if the transcriptional profile of this cluster results from the presence of one or multiple distinct cell types, these might be at risk of being lost due to technical limitations of scRNA-seq experiments of the liver. Cluster 5 spots show expression of mesenchymal cell makers and are associated with “collagen fibril organization” pathways. We propose that cluster 5 might represent parts of the Glisson’s capsule, composed of collagen fibrils together with its underlying mesothelium, representing the connective tissue encapsulating the liver and regions with thicker, hilar periportal mesenchyme. The capsule preserves the structural integrity of the loosely constructed liver and enables the division into lobes ^47^.

The mesenchymal cell-marker *Vim* is reported to maintain mesenchymal cell structure and serves as an indicator for cell proliferative activity in liver cells ^25,54^ and *Gsn*, encoding the actin-binding protein GELSOLIN has an anti-apoptotic role in the liver ^55^. Anti-apoptotic effects and enrichment of connective tissue, possibly from the Glisson’s capsule, might be crucial in fragile positions of the organ or close to connection positions of liver lobes. The two additional pathways involved in structural integrity in cluster 5, namely “extracellular matrix organization” and “extracellular structure organization”, further advocate for a structural function of cells in this cluster. Enrichment of gene ontologies associated with response to cytokines are observed in, but not limited to, cluster 5; hence they are contributing rather than defining components of the cluster’s expression profile and function of the structure.

Knowledge about the functional reason for division of the liver into multiple lobes, and the maintenance of structural integrity in the organ, is still incomplete ^19,56^. Considering the sample size used in this study, we can provide first indications rather than general claims of the function of this proposed structure. On top of capturing and supporting previously observed trends of tissue heterogeneity in the mammalian liver, our study serves as a valuable resource to further investigate the spatial expression of structural components and gene candidates involved in the aforementioned processes.

In summary, this study presents a novel approach to investigate the transcriptional landscape of liver tissue through spatial transcriptomics and innovative computational approaches. We designed and implemented computational tools allowing physical distance measurements and predictions of veins. In addition, we are proposing the presence of transcriptionally distinct structures in liver tissues that previously have not been reported using transcriptomic analyses, likely due to the rarity of cells contributing to these structures.

With anticipated future advances in the spatial genomics field, increased resolution will promote detailed investigations of rare cell types in tissue space. This study constitutes a compelling initial exploration of the benefits that spatial transcriptomics provides for studies of the liver and consider it a valuable data resource for the hepatology field. We further anticipate that ST will be highly beneficial for future studies addressing liver development, immunity and general pathology.

## METHODS

### Ethical statement

The Regional Animal Research Ethical Board, Stockholm, Sweden, approved experimental procedures and protocols involving extraction of organs from mice (N135/15, N78/16 and 9707-2018), following proceedings described in EU legislation (Council Directive 2010/63/EU).

### Total RNA extraction

To test for the RNA quality of the tissue for further downstream analysis, the tissue was sectioned and up to 8 sections of 10 µm sections were placed in Lysing Matrix D tubes (MPBiomedicals, cat.no.: 116913050-CF) containing Buffer RLT Plus (Quiagen, cat.no.: 1053393) and ß-Mercaptoethanol (Thermofisher, cat.no.: 31350010) and homogenized in a Fastprep instrument (ThermoSavant). The flowthrough was collected through a gDNA Eliminator column and 250 ul of pure ethanol was added. Total RNA was further extracted using the RNAeasy mini kit (Quiagen, cat.no.: 74104) according to manufacturer’s instructions. The RNA integrity number (RIN) for each sample was assessed performing Bioanalyzer High Sensitivity RNA Analysis (Agilent cat.no.: 5067-1535).

### Collection and preparation of liver samples

Female C57BL/6 mice (Charles River), housed under specific pathogen-free conditions at the Experimental Core Facility, Stockholm University,were euthanized between week 8 and 12, livers were collected, and four lobes were separated. Each lobe was segmented so cryosections would fit on the 6,200 x 6,400 µm areas of the Codelink-activated microscope slides and frozen in −30°C 2-Methylbutane (Merck, cat.no.: M32631-1L). The frozen liver samples were embedded in cryomolds (10×10 mm, TissueTek) filled with pre-chilled (4°C) OCT embedding matrix and frozen (CellPath, cat.no.: 00411243). For downstream experiments, the frozen samples were sectioned at 10 µm thickness with a cryostat (Cryostar NX70, ThermoFisher). Each subarray on the slide is covered with 1934 spots with a 100 µm diameter, containing approximately 200 million uniquely barcoded oligonucleotides with poly-T_20_VN capture regions per spot (Barcoded slides were manufactured by 10X Genomics Inc, probes were manufactured by IDT). The full protocol, including sequencing and computational analysis was performed for 8 sections across 3 samples. Sample 1 and sample 3 each include 3 sections of the caudate lobe. Sample 2 includes 2 sections of a liver piece of the right liver lobe. All sections of all samples have undergone the same treatment.

### Histological staining and annotations

We performed the spatial transcriptomics workflow according to Ståhl et al. and Vickovic et al., respectively ^57,58^. After 10 minutes of formalin fixation of the tissue on the slides they were dried with isopropanol and stained with Mayer’s hematoxylin (Dako, cat.no.: S330930-2) and bluing buffer (Dako, cat.no.: CS70230-2) followed by Eosin (Sigma-Aldrich, cat.no.: HT110216-500ML), diluted in Tris/acetic acid (pH 6.0). The stained sections were mounted with 85% glycerol (Merck Millipore, cat.no.: 8187091000) and covered with a coverslip. Bright field images were acquired at 20x magnification, using Zeiss AxioImager 2Z microscope and the Metafer Slide Scanning System (Metasystems). The liver images were assessed by a liver expert (NVH) who annotated the portal (blue) and central (red) veins, based on the presence of bile ducts and portal vein mesenchyme (PV) or lack thereof (CV). When the quality of the sample did not allow for annotation, “ambiguous vein” (green) was reported.

### Permeabilization, cDNA synthesis, tissue removal and probe release

Next, the slides were put in mask holders (ArrayIT) to enable separated on-array reactions in each chamber as described previously ^58^. Each tissue section was pre-permeabilized using Collagenase I for 20 minutes at 37°C. Permeabilization was performed using 0.1% pepsin in 0.1 M HCl for 10 minutes at 37°C. cDNA synthesis was performed overnight at 42°C. Tissue removal from the arrays prior to probe release was performed using Proteinase K in PKD buffer at a 1:7 ratio at 56°C for 1 hour. Lastly, the surface probes were released and cDNA library preparation followed by sequencing was performed.

### cDNA library preparation and sequencing

Released mRNA-DNA hybrids were further processed to generate cDNA libraries for sequencing. The sequencing libraries were prepared as described in Jemt *et al*. ^59^. In short, the 2^nd^ strand synthesis, cDNA purification, in vitro transcription, amplified RNA purification, adapter ligation, post-ligation purification, a second 2^nd^ strand synthesis and purification were done using an automated MBS 8000+ system. To determine the number of PCR cycles needed for optimal indexing conditions, using qPCR as described previously ^58^. After determination of the optimal cycle number for each sample, the remaining cDNA was indexed and amplified. The indexed libraries were then purified using an automated system as previously described ^60^. The average length of the indexed cDNA libraries was determined with a 2100 Bioanalyzer using the Bioanalyzer High Sensitivity DNA kit (Agilent, cat.no.:5067-4626), concentrations were measured using a Qubit dsDNA HS Assay Kit (Thermofisher, cat.no:Q32851) and libraries were diluted to 4nM. Paired-end sequencing was performed on the Illumina NextSeq500 platform, with 31 bases from read 1 and 46 bases from read 2 resulting in the generation between 15 and 32.1 million raw reads per sample. To assess the quality of the reads fastqc (v 0.11.8) reports were generated for all samples.

### Spot visualization and image alignment

The staining, visualization and imaging acquisition of spots printed on the ST slides were performed as previously described ^57^. Briefly, spots were hybridized with fluorescently labeled probes for staining and subsequently imaged on the Metafer Slide Scanning system, similar to the previous acquisition of the HE images. The previously obtained brightfield of the tissue slides and the fluorescent spot image were then loaded in the web-based ST Spot Detector tool ^61^. Using the tool, the images were aligned and the spots under the tissue were recognized by the built-in recognition tool. Spots under the tissue were slightly adjusted and spots under the tissue were extracted.

## Computational analysis

### Mapping, gene counting and demultiplexing

Processing of raw reads was performed using the open source ST Pipeline (v 1.7.6) ^62^. In short, quality trimming was performed and homopolymer stretches longer than 15 bp were removed. The reads were subsequently mapped to the annotated reference genome (GRCm38 v86 and corresponding GENCODE annotation file) using STAR (v 2.6.1e) ^63^. The trimmed and filtered reads were demultiplexed according to their respective 18 nucleotides spatial barcode using TagGD demultiplexing ^64^ . After filtering, PCR duplicates were removed and gene count matrices were generated.

### Dimensionality reduction and clustering

Main computational analysis of spatial read-count matrices was performed using the STUtility package (v 0.1.0) ^65^ in R (v 4.0.2). The complete R workflow can be assessed and reproduced in R markdown (see code availability section). First, count matrices and metadata were loaded, translating Ensembl IDs to gene symbols simultaneously. Reads of individual samples were filtered to keep only protein-coding genes and subsequently normalized using the “SCTransform” function in Seurat. The created objects were then integrated using the canonical correlation analysis (CCA) with the MultiCCA function provided in https://github.com/almaan/ST-mLiver.

Normalization of integrated data was performed, regressing out sample identities using the “SCTransform” function in Seurat. Thereafter, the CCA vectors were subjected to shared-nearest-neighbor (SNN) inspired graph-based clustering via the “FindNeighbors” and “FindClusters” functions. For modularity optimization, the louvain algorithm was used and clustering was performed at a resolution of 0.3 for clustering granularity.

### Visualization and spatial annotation of clusters

To visualize the clusters in low-dimensional space and on the spot coordinates under the tissue, non-linear dimensionality reduction was performed using UMAP with the CCA vectors as input. Visualization and annotation of identified clusters in UMAP space, on spot coordinates as well as superimposed on the Hematoxylin- and Eosin images was performed using the Seurat and STUtility package.

### Differential gene expression analysis (DGEA) and expression programs

Differential gene expression analysis of genes in identified clusters was performed using the function “FindAllMarkers” from the Seurat package. Following the default option of the method, differentially expressed genes for each cluster were identified using the non-parametric Wilcoxon rank sum test. Initial thresholds were set to a logarithmic fold-change of 0.25 to be considered differentially expressed in a cluster and to be present in at least 10% of the spots belonging to the same cluster. Representative markers for each cluster were further selected, by choosing genes with a positive logarithmic threshold above 0.5 and an adjusted p-value below 0.05. P-value adjustments are based on bonferroni correction using all genes in the dataset.

After the identification of marker genes of the individual clusters, we identified expression programs of genes for clusters we identified to have spatial distribution in our data. These were cluster 1 (periportal cluster), cluster 2 (pericentral cluster) and cluster 5. Creation of expression programs was performed using the “AddModuleScore” function in Seurat. In brief, we stored the marker genes of each cluster in a list to serve as input for the function. From this input, the average expression of each program (list of markers) was calculated for each spot under the tissue and subtracted by the aggregated expression of a control gene set. Here the control gene set included all genes present in our data. All analyzed genes were then binned based on averaged expression and with the default number of 24 bins for the function, and 100 control genes of the control feature set were randomly selected from each bin. Higher scores indicate more marker genes of the program to be highly expressed in a spot, while lower scores indicate that no or only a small number of genes is expressed at low levels in the spot.

### Spatial autocorrelation

To explore the correlation between spatial distribution and expression of all genes in our data we performed spatial autocorrelations using the CorSpatialGenes of the STUtility package. The method is based on building a connection network from the spot-coordinates for each spot and the four surrounding neighbors at a maximum distance of 150µm. Thereafter, individual connection networks are combined to a tissue-wide connection network to compute autocorrelations for the whole dataset. Based on the neighbor groups of each spot, lag vectors for all input features are calculated, essentially being the sum expression of the respective feature in the neighbor spots. This considered, neighbouring spots with high spatial autocorrelation of features demonstrate similar expression levels. This allowed us to compute the correlation score between the lag vector and the actual expression vector to estimate spatial autocorrelations.

### scRNA-seq data

Publicly available scRNA-seq data was analyzed to compare and complement the spatial data in our studies. Two datasets were downloaded for this purpose, the scRNA-seq data set of cells originating from liver tissue from the Mouse Cell Atlas (accessed 2020-10-06) ^39^ and differential gene expression data from the single cell spatial reconstruction of mouse liver ^14^. For comparative analysis and visualization, scRNA-seq data of the Mouse Cell Atlas was analyzed using the Seurat package (v 3.2.2). The count-data was first filtered for mitochondrial genes and normalized using the “SCTransform” function. Dimensionality reduction was performed using PCA and graph-based clustering was performed using the “FindNeighbors” and “FindClusters” function with a resolution of 0.8 for clustering granularity. Visualization of the clusters in low-dimensional space was performed using non-linear dimensionality reduction (UMAP). Clusters were grouped by the cell type annotations provided by the metadata of the single cell data set. The second data-set used for comparative analysis was extracted from single cell spatial reconstruction data ^14^. Differential gene expression data between layers of zonation was compared to markers for pericentral or periportal zonation in our data set using R (v 4.0.2).

### Correlation analysis

Correlation analyses between genes of clusters were performed using Pearson correlation, establishing linear correlations between differentially expressed genes of the clusters in base R. Visualization of correlation values was carried out using the corrplot package (v 0.84). The correlation coefficients of the matrix were ordered using the method “FPC”, describing the first principal component order of the correlation coefficients.

To explore the correlation relationship between single cells (assigned to the classes “pericentral hepatocytes” and “periportal hepatocytes”) and the spatial transcriptomics “pericentral (cluster2)” and “periportal (cluster1)” clusters, spearman rank correlation coefficients were calculated. First module scores of genes assigned to each cluster were calculated for each set of data: spatial transcriptomics and single cell data of the Mouse Cell Atlas. Notably, not all genes present in one data-set were present in the other, therefore only genes present in the respective dataset were considered. Thereafter, spearman rank correlation between the scores for all groups (“pericentral hepatocytes”, “periportal (cluster1)”, “pericentral (cluster2)”) were performed. The relationships were visualized using the corrplot package, with values ordered in the original input order.

### Pathway analysis

Functional enrichment analysis of marker genes of clusters was performed using gprofiler2 (v 0.1.0). For the analysis we extracted the gene symbols of each cluster and stored them in a list. The function “gost” of the gprofiler2 package was then used to perform gene set enrichment analysis, on input marker gene lists. In short, the function maps genes to known functional information sources and detects statistically significantly enriched terms. Since our data consists of murine liver sections, the organism was set to *mus musculus* and the source was set to Gene Ontology (GO) biological processes. Visualization of the 5 most significantly enriched processes for cluster 5 was performed using ggplot2 (v 3.3.2). Significance was adjusted using g:SCS (Set Counts and Sizes), as originally described by the authors of the gprofiler package ^66^. Enrichment scores are represented as the negative log10 algorithm of the corrected p-value. For visualization of functional enrichment on the tissue coordinates, marker genes of cluster 5 were referenced against all genes belonging to the go-terms for “collagen fibril organization (GO_0030199)” and “response to cytokine (GO_0034097)”, extracted from the gene ontology browser of the Mouse Genome Informatics database. Gene expression programs were generated for genes belonging to each gene ontology term as described before and visualized on the spots.

### Single cell data integration (stereoscope)

The spatial data was integrated with the MCA dataset using *stereoscope*, a probabilistic method designed for spatial mapping of cell types ^38^. In short, *stereoscope* models both single cell and spatial data as negative binomial distributed, learns the cell type specific parameters from the (annotated) scRNA-seq data, and uses these to deconvolve the gene expression in each spot into proportion values associated with respective cell type. *stereoscope* uses a stochastic gradient descent approach, leveraging the PyTorch framework, to obtain the maximum likelihood/maximum *a priori* estimates of both the parameter estimates and proportion values. In both steps (parameter estimation and proportion inference) a batch size of 2048 and 50000 epochs were used, a custom list of highly variable genes - see next section for details - was used rather than the full expression profiles; default values were used for all other parameters. *stereoscope* can be accessed at https://github.com/almaan/stereoscope, where more detailed documentation regarding the parameter values is provided. The *stereoscope* version used in the study was v.0.3 (commit: aacd5f775b73b138e504c35ff0cb3ffafbfc78ff)

The cell type proportion values were overlaid on the tissue section images by using the “FeatureOverlay” function in the STUtility package. To make our visualization more robust to outliers, we scaled all the proportion values using what we refer to as *quantile scaling*. Here, this procedure was performed in two steps: First, all values larger than the 0.95 quantile are changed to this quantile value (i.e., the data is clipped); then, within every cell type and section we divide the clipped values by their maximum, effectively mapping them to the unit interval [0,1]. Thereafter, the proportion values for all 20 cell types in the single cell dataset were plotted on the spot coordinated and overlaid on the Hematoxylin and Eosin stained tissue sections.

### Selection of highly variable genes for *stereoscope*

Seurat (v 3.2.2) was used to extract a set of highly variable genes from the MCA single cell data, following the procedure recommended in the online Seurat *Clustering Tutorial* (https://satijalab.org/seurat/v3.2/pbmc3k_tutorial.html). To elaborate, the following two steps were applied to the MCA single cell data set in sequential order: data normalization (*NormalizeData*, default parameters), and identification of highly variable genes (*FindVariableFeatures, selection*.*method = “vst”, features = 5000*). The complete set of extracted genes was used in the *stereoscope* analysis (Supplementary table 7).

### Cluster Interaction Analysis

To gauge the extent to which the expression-based clusters interacted in the spatial data, we constructed a simple interaction analysis based on a nearest neighbor approach. First, for every spot within each cluster, the cluster identity of the four nearest neighbors within a distance threshold are registered. To avoid confusion, note how these distances refer to separation of spots in the ST array and not in gene expression-space. The distance threshold is used to ensure that only spots in the actual physical neighborhood are included in the count, as might not be the case for spots near the edge otherwise. Second, once neighbor identities have been registered for all members of a cluster, we convert these integer values to a fraction by dividing them with the total number of neighbors associated with the cluster. Thus, for any given cluster we have a set of *n_cluster* values representing the total fraction of neighbors that belong to each one of the clusters. Since spots need to be positioned somewhere in space, clusters with a large member count will by default neighbor more spots than a cluster with low member count. Hence, to assess whether an interaction seems to be present or not, one must account for cluster size and spatial organization of the spots; here done by random permutation of the cluster labels (100 times) followed by re-calculation of the same neighborhood fraction values. This allows us to put the observed neighborhood fractions into context, to what might be expected by random chance given the cluster cardinalities and spot organization. It is by this approach that Supplementary Figure 2 was generated, where each bar represents the observed values, the dashed black line the empirical mean value from the permutation analysis, and the magenta envelopes filling the area of two standard deviations from the mean.

### Features as a function of distance

To examine how certain features of interest (e.g., gene expression or proportion values) were influenced by the physical proximity to morphological structures (e.g., central and portal veins) in the tissue samples, an approach to model these values as a function of the distance to said structure was devised. This procedure is described in detail below:

Using the brightfield HE-images, a mask was created for each morphological structure. These masks covered all pixels considered to belong to the structure. Each structure was assigned an individual (numerical) id, and one or more class attributes related to it (e.g., “*vein type*”). As the spots’ (capture locations) positions relate to pixel coordinates in the HE-image, it was possible to - computationally - measure the distance from a spot to each of these structures.

The distance (*d(s,t)*)) from a spot *s* to a structure *t* was here defined as *the minimal (euclidean) distance from the center of spot s to any pixel p belonging to the mask of t”*. In other words, if *M_t_*is the set of all pixels in the mask belonging to structure *t* then:

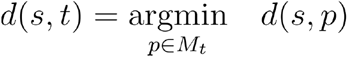

The same procedure was used when determining the distance to a specific class attribute (e.g., vein type), except that the *union of all masks associated with a structure of said class* was used instead of only a single mask. That is, if *M_C_* is the set of all pixels belonging to any structure of class *C*, then the distance (*d(s,C))* between spot *s* and class *C* is:

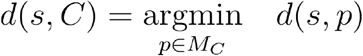

Next, once distances were determined, for a feature *x* of interest (e.g., expression value) and a structure *t*, a tuple (*d(s,t),x_S_*) was formed for each spot *s*; i.e. the distance for every spot was associated with the value of the feature. This set of distance-feature tuples could then be visualized in graphs, in order to depict the feature values’ dependence on their distance to the structure.

To better capture general trends in the data, *scatterplot smoothing* (using the *loess* function from the scikit-misc package, v 0.1.3, default values for all parameters), was applied to generate smoothed estimates. The smoothed values would then serve as an approximation of a function *f* such that x_s_= *f(d(s,t))*, to be interpreted as if the feature value is a function of the distance to the structure. Plotting the smoothed values against their associated distances results in a visualization of the function approximation over the distance domain, these plots are referred to as “*feature by distance”* plots; where the feature for example could be *expression* or *proportion*.

To better account for synergies between structures of various classes, a variant of the feature by distance plots was implemented. Rather than considering features as a function of the distances between spots and structures of *one* specific class *C* (*d(s,C)*), the logged ratio between distances to *two* classes of interest (e.g., *C_1_*and *C_2_)* was used, more specifically:

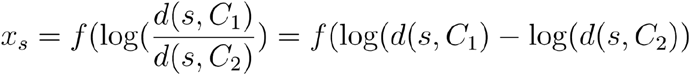

Negative values of the log-ratio represent positions in the domain (the tissue) where the distance to the second class (C_2_) is larger than to the first class (C_1_) Positive log-ratio values have the opposite interpretation. The same smoothing procedure as described above was used to approximate this function. Plotting the smoothed values against their associated log-ratio values produces the visualizations referred to as “*feature by distance-ratio”* plots. The two classes used in this study were *central veins* and *portal veins*, but the concept is generalizable to any pair of classes.

Unless otherwise stated, feature-distance tuples across all sections were aggregated when generating features by distance/distance-ratio plots. The envelopes encapsulating the smoothed approximation represent one standard error (SE) as given by the loess algorithm.

### Expression-based classifier

To assess whether the gene expression of a structure’s (e.g., central or portal vein) neighborhood held sufficient information to infer its class, we constructed a classifier designed to predict structure-class based on gene expression data. The steps of data processing and explicit details for the classification procedure are described below:

First, *neighborhood expression profiles* (NEPs) for each structure were created, representing a weighted (by distance) average expression of a set of features (here genes) in the neighborhood of a structure. The neighborhood (*N(t)*) of a structure *t* was defined as the set spots with a distance less than a threshold *T_N_* to *t*. That is:

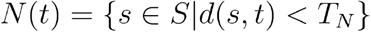

Where *S* is the set of all spots, while distances between spots and structures (d(s,t)) are defined and computed as described in the section above (*Features as a function of distance*). In this study, we set the distance threshold (T_N_ to 210). Having formed the neighborhoods, their associated expression profiles for a feature (*x_N(t)_*) were assembled accordingly:

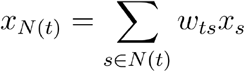

Where *w_ts_* are the distance-based weights given by:

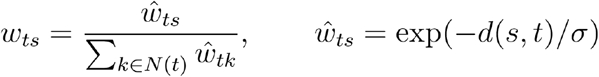

In this analysis, σ was set to 20. As multiple features (*x*) are used, NEPs are represented by a vector of *N* (the number of features used) elements, denoted as ***x_N(t)_***. Each NEP was then given a class label, *portal* or *central*, based on the associated structure’s annotations. The task of predicting class labels from the NEPs then surmounts to a *multivariate binary classification problem*, for which a logistic regression model was employed. Implementation-wise the logistic regression was performed by using the *LogisticRegression* class from *sklearn’s* (v 0.23.1) *linear_model* module, a *l2* penalty was used (regularization strength 1), the number of max iterations was set to 1000, default values were used for all other parameters.

In short, the logistic model considers the class label (*z_t_)* of a structure *t* as bernoulli variable conditioned on the NEPs, i.e:

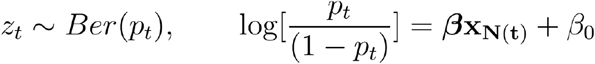

Fitting the model equates to finding the maximum likelihood estimates of the **β_N(t)_** and β_0_given the observed data and regularization terms. Once fitted, the class of a structure *t* is taken as class 1 if p_t_≤ 0.5 and class 2 if p_t_> 0.5. However, p_t_ is a continuous value that also can be interpreted as the probability of a structure belonging to each class (low values indicate more similarities with class 1 and vice versa) - offering a form of *soft* classification.

To validate performance, cross validation strategies were implemented at two different levels: *section* and *sample*. In the former K-sections were set aside forming a *test set* while the remaining sections constituted the *training set*. The model was trained on the *training set* and evaluated on the K sections in the test set. This procedure was iterated for all combinations of the pairs. Cross validation on the sample level was conducted in a similar fashion, but setting w.r.t. samples rather than individual sections - here setting aside a sample is equivalent to excluding *all sections associated with the sample*.

As the number of samples - and thus structures - were fairly low, using the complete expression profiles (i.e., all genes) would likely have led to an overfitted model (n_features >> n_samples). Thus, a reduced set of genes were used to construct the NEPs, extracted from the set of marker genes identified in the previously described differential gene expression analysis - this set of genes can be found in Supplementary figure 13.

## Data Availability

The datasets generated during and/or analyzed during the current study are available in the doi-minting repository ST Liver (10.5281/zenodo.4399655) and can be accessed at https://zenodo.org/record/4399655. The data used for comparative analysis of previously published data can be accessed at *14* and Gene Expression omnibus (accession code GSE84498) as well as at ^39^, http://bis.zju.edu.cn/MCA/ with raw data accessible at Gene Expression omnibus (accession code GSE108097)

## Code Availability

Code to reproduce the analysis can be accessed at https://github.com/almaan/ST-mLiver, it has also been deposited to a doi-minting repository (Zenodo) accessible via https://zenodo.org/record/4399655. Functions and classes pertaining to the *feature by distance* and *classification* analysis have been assembled into a Python module (*hepaquery*), while the workflow used to produce the results is given in a set of notebooks. A CLI program to prepare the data for the distance-related analysis once masks have been created is also provided. See the repository documentation for more information regarding reproduction of the analyses.

## ACKNOWLEDGEMENTS

We would like to thank Carina Olivera and Emily C. Ross, for providing excellent technical support and expertise in the animal work for this study.

This research project was generously funded by grants from: the Swedish Society for Medical Research (SSMF), The Swedish Research Council (VR) and The Jeansson Foundation to J.A; The Swedish Research Council (VR) to J.L.; The Swedish Research Council (VR), the Swedish Foundations Starting Grant/Ragnar Söderberg Foundation, and Karolinska Institutet to E.R.A.; The Sven and Lily Lawski Foundation to F.H., and The European Association for the Study of the Liver/EASL to J.M.

## AUTHOR CONTRIBUTIONS

**J**.**A**. conceived the study; **J**.**L**. and **J**.**A**. supervised the project; **F**.**H**., **A**.**M**. and **S**.**K**. optimized the ST methodology for liver; **F**.**H**. and **S**.**S**., designed and carried out ST experiments; **F**.**H**., **L**.**L**. and **A**.**A**. developed computational methodology and analyzed data; **L**.**L**. generated CCA function; **F**.**H**. and **A**.**A**. designed and implemented the expression by distance and cluster distribution analysis; **N**.**V**.**H**. performed histological annotations; **E**.**R**.**A**., **N**.**V**.**H**., **J**.**M**. provided essential background on liver biology; **A**.**B**., **E**.**R**.**A**., **N**.**V**.**H**., **J**.**M**. and **E**.**E**. provided technical aspects on liver assays; **F**.**H**. and **J**.**A**. wrote the manuscript with the help of **A**.**A**., **E**.**R**.**A**. and **J**.**L**. All authors edited and gave critical input on the manuscript.

## Additional information

### Supplementary information

is available for this article. Correspondence and requests for materials should be addressed to

