## Supplementary Figures 1-13, Supplementary Table 4 for "Spatial Transcriptomics to define transcriptional patterns of zonation and structural components in the liver"

**Supplementary figure 1:** Visualization of unsupervised clustering results of integrated sequencing data on spots under the tissue for **a)** replicate 1 of three sections of a part of a caudate lobe **b)** replicate 2 of three sections of a part of a caudate lobe and **c)** replicate 3 of two sections of a part of a right lobe.

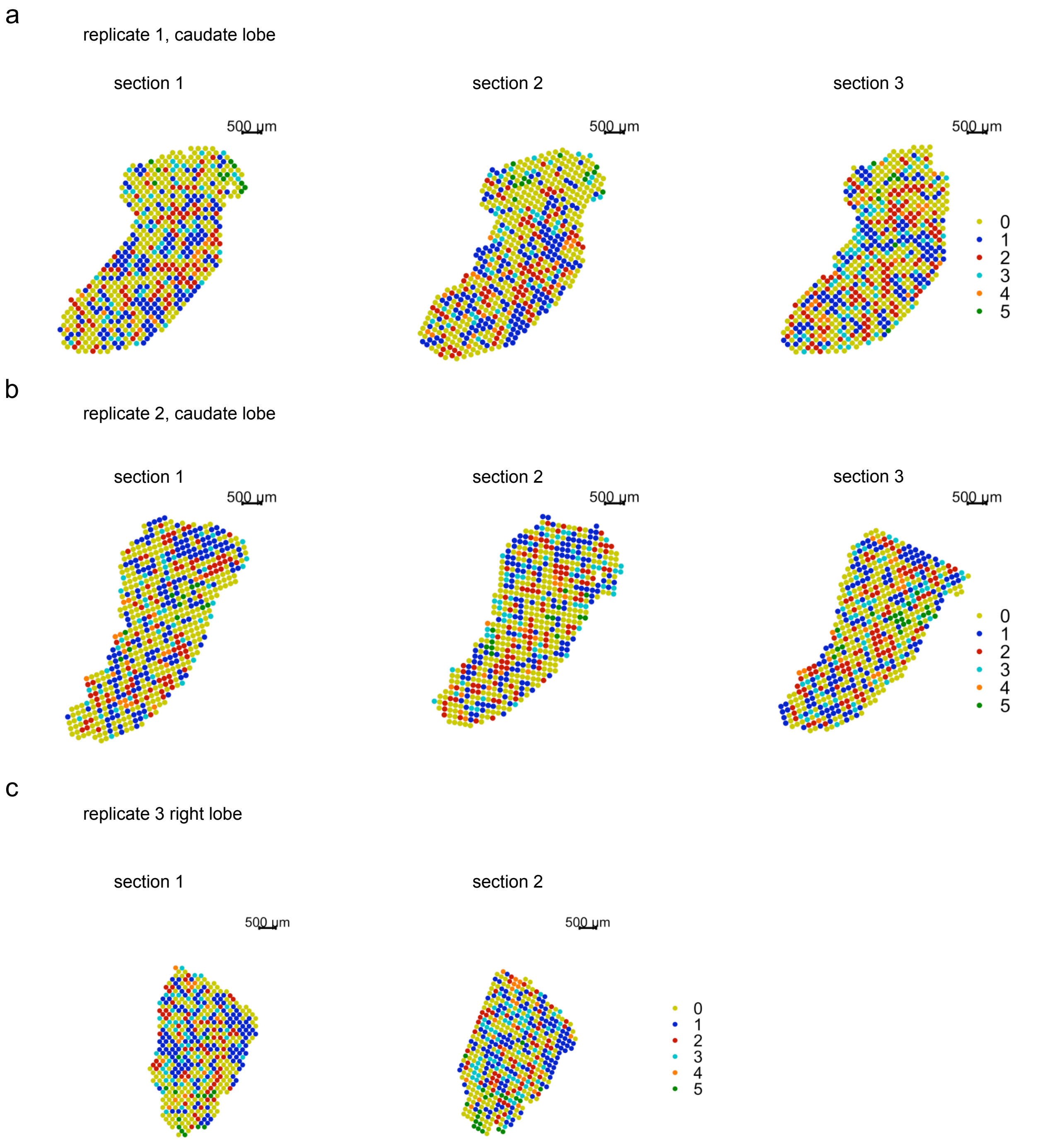

**Supplementary figure 2:** For spots belonging to each cluster identity (0-5) the most frequent cluster identities of the four spots within the closest proximity (150 μm) were determined, ranging from fractions between 0 and 0.5 of all neighboring spots of each cluster. The cluster for which the neighboring fractions are determined is depicted in grey with a red dashed stroke. Dashed lines in magenta indicate the expected distribution of fractions among the neighboring clusters if the spots under the tissue were randomly assorted.

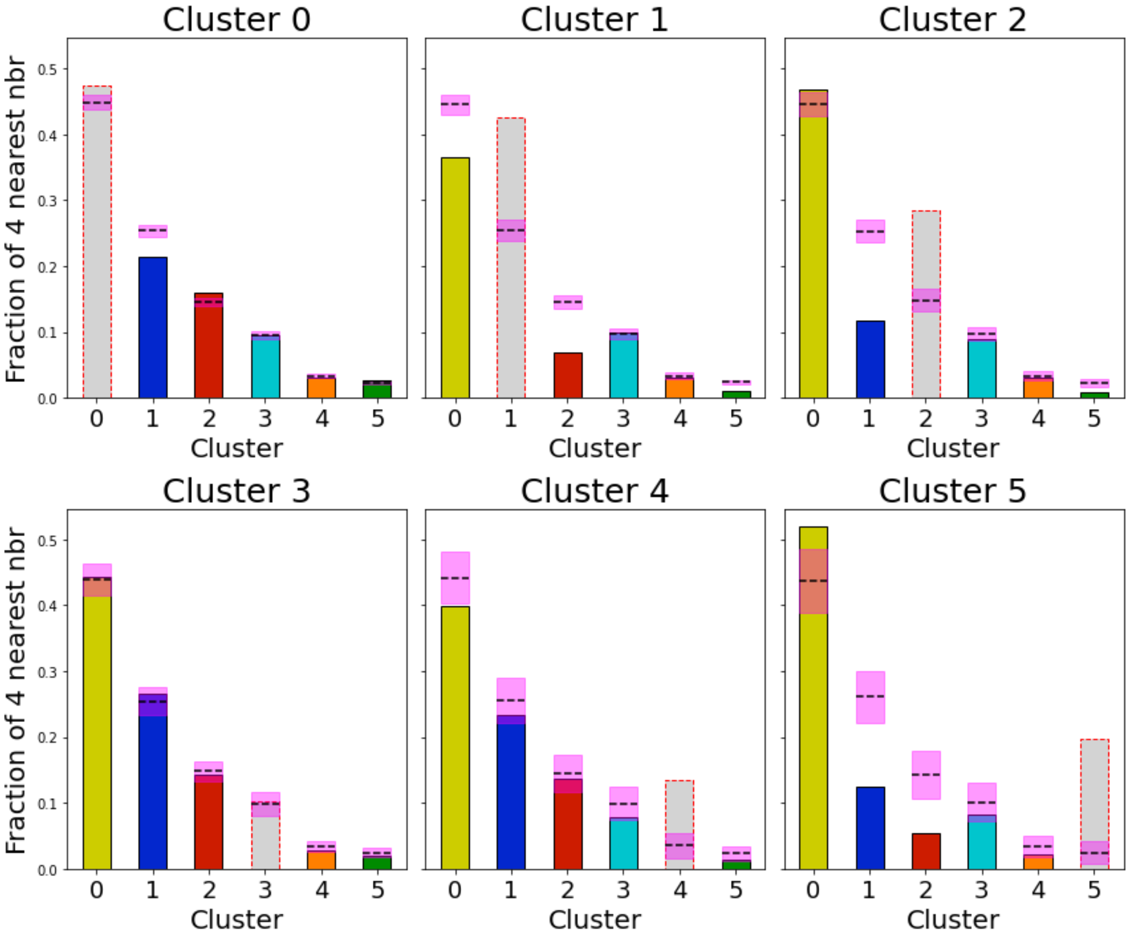

**Supplementary figure 3.1 - 3.3:** Visualization of single cell data integration by *stereoscope* on spots under the tissue for all sections of this study. Spot-wise calculated proportion-values were scaled using quantile scaling between values of 0 to 1 for all 20 annotated cell types of the MCA single-cell data and each section. Proportion-values are visualized on spots under the tissue for each experiment . **3.1** displays section1 (top) and section 2 (bottom) of the a right lobe. **3.2** displays section 1 (top) to section 3 (bottom) of a caudate lobe. **3.3** displays section 1 (top) and section 3 (bottom) of a caudate lobe of a second experiment.

#### Supplementary figure 3.1

replicate 3, right lobe

section 1

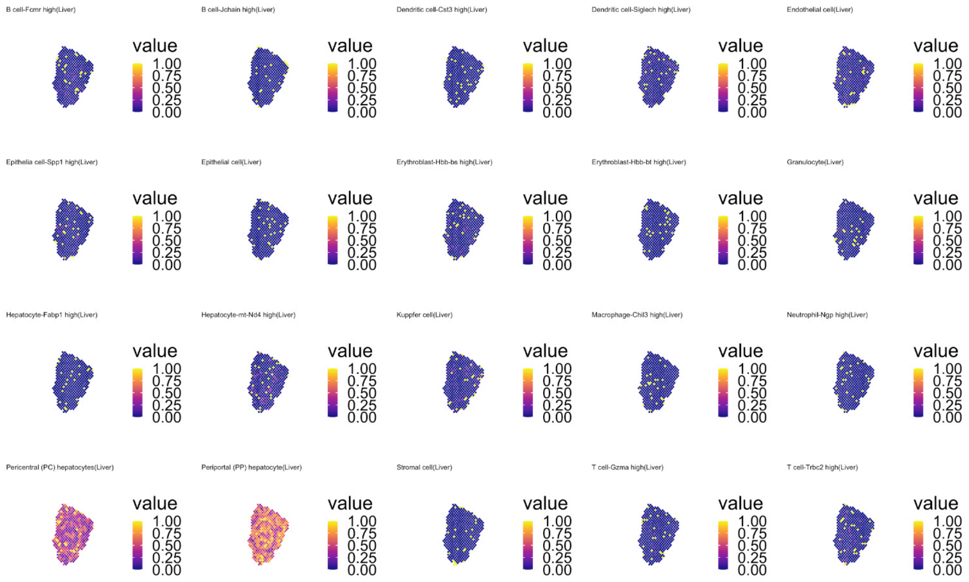


section 2

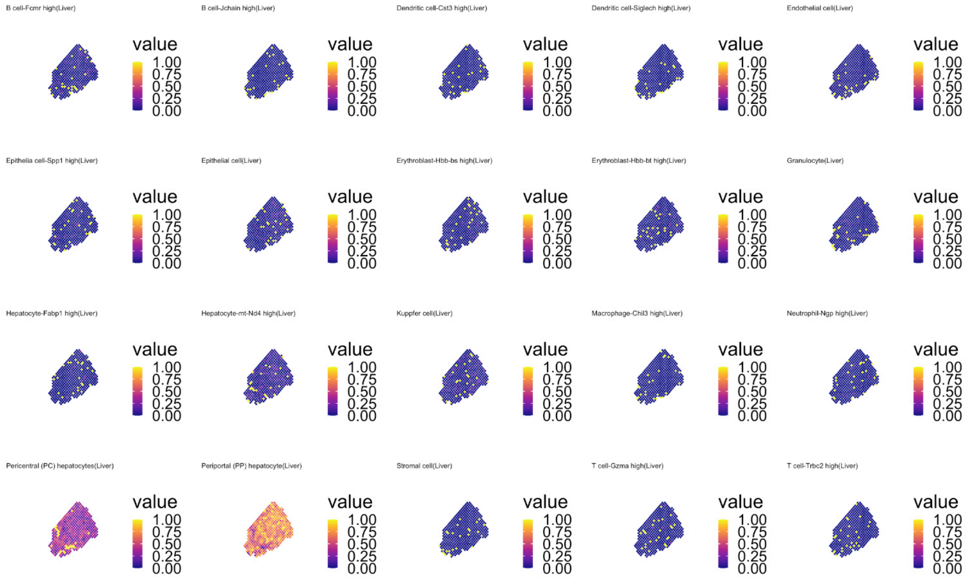

### Supplementary figure 3.2

replicate 1, caudate lobe

section 1

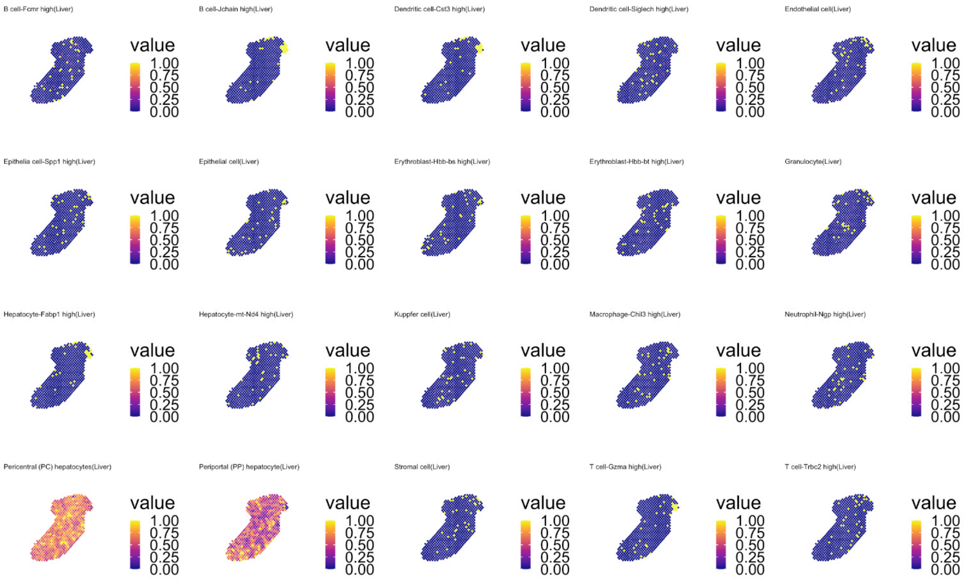


section 2

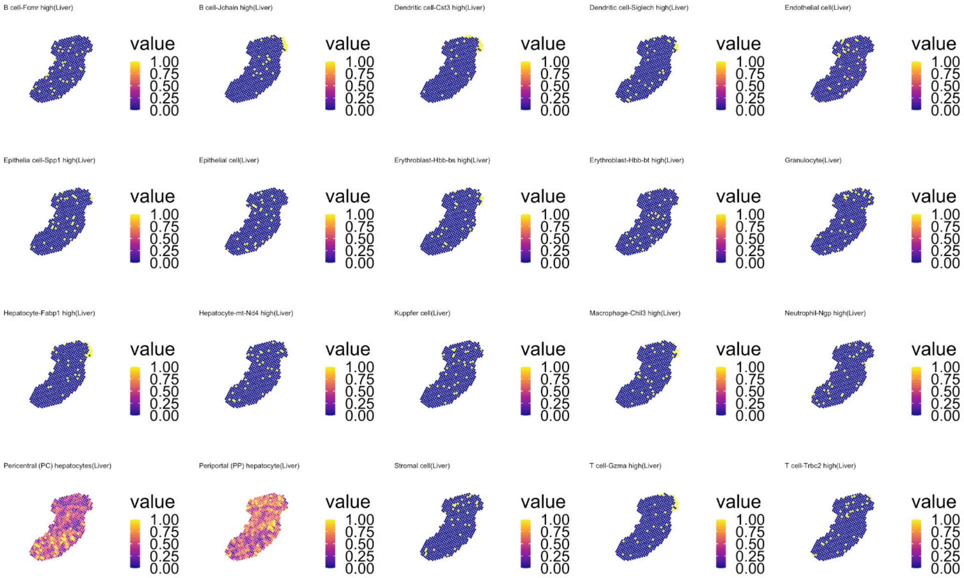


section 3

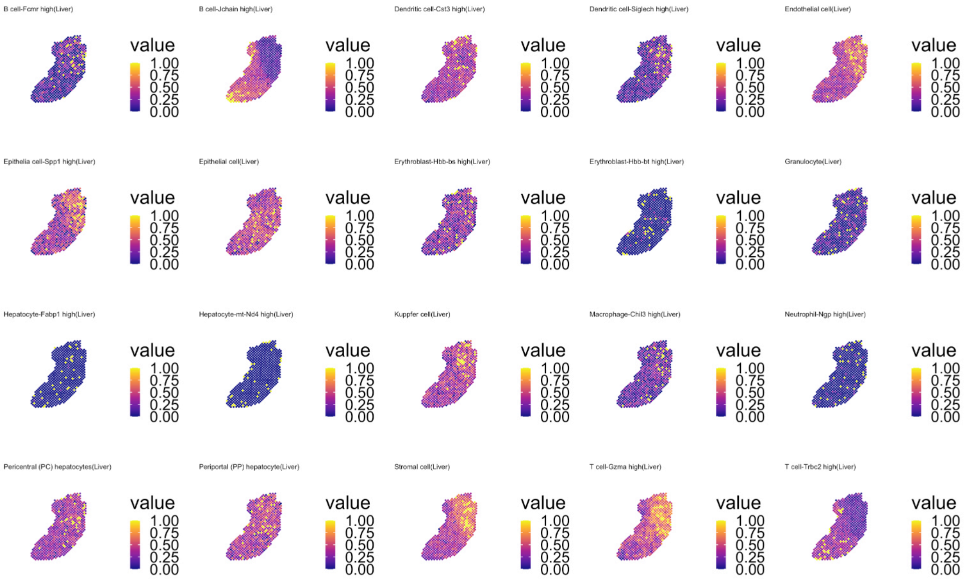

### Supplementary figure 3.3

#### replicate 2, caudate lobe

##### section 1

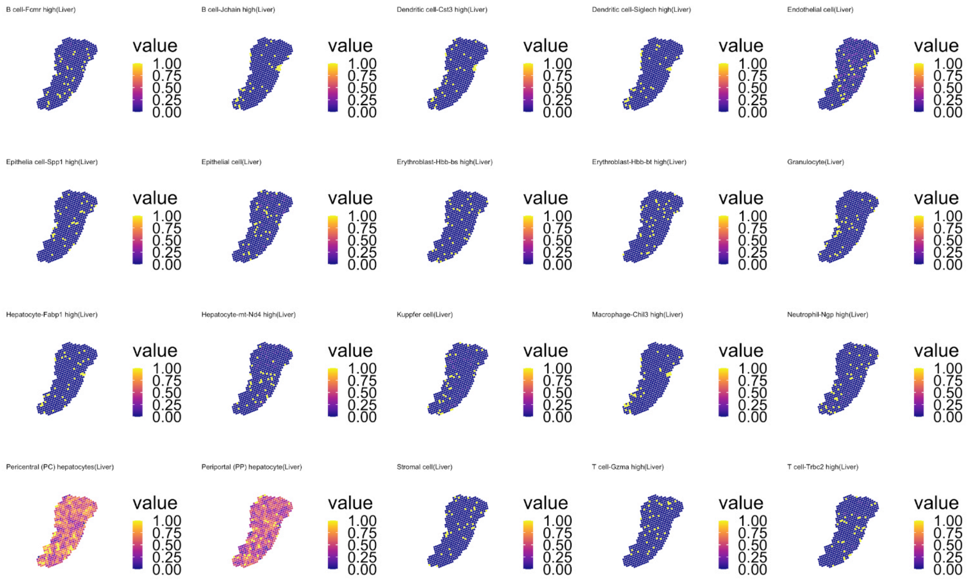


##### section 2

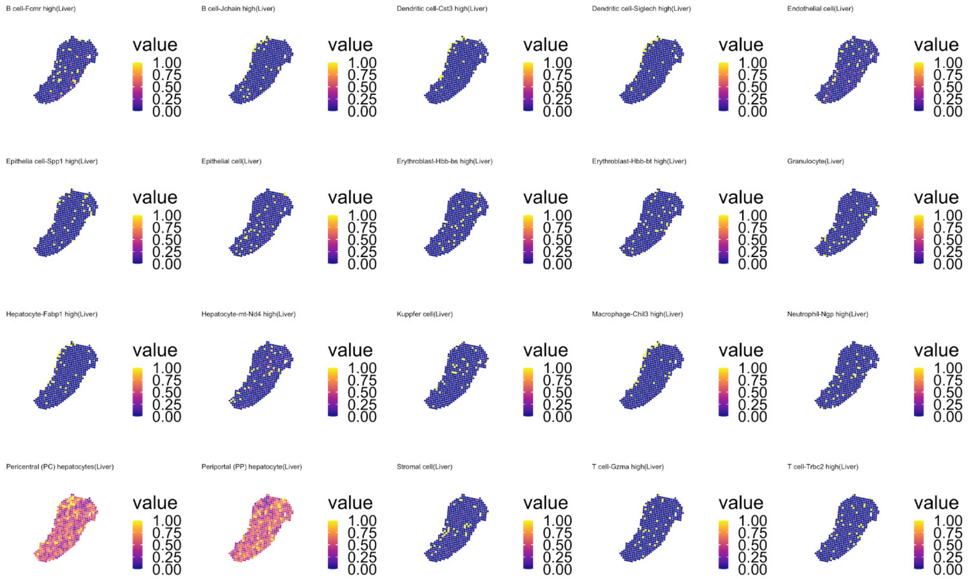


##### section 3

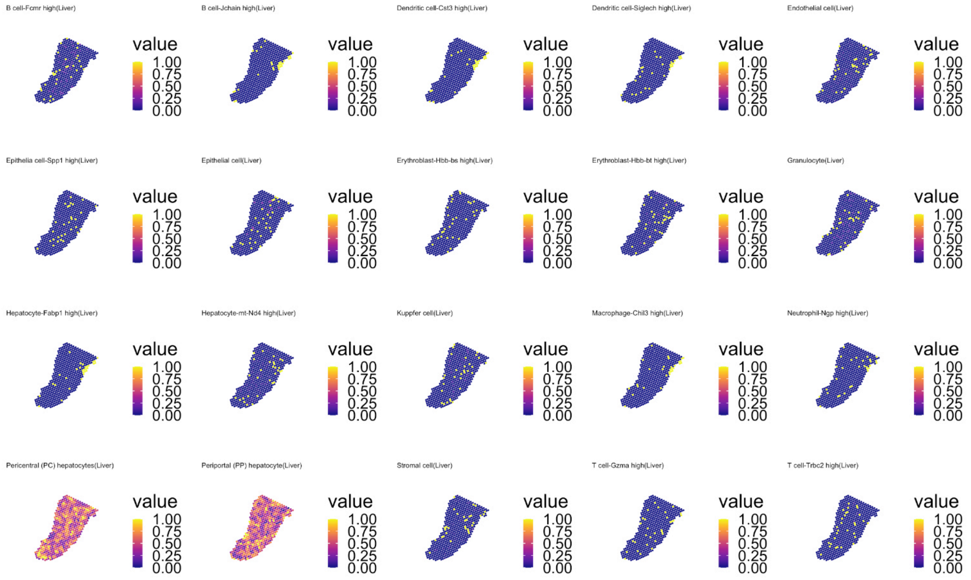

**Supplementary figure 4:** Spearman correlations between single cell periportal and pericentral cell-types and pericentral and periportally annotated spots in spatial transcriptomics data.
**a)** Spearman correlations between module score values of markers from spatial transcriptomics data (cluster 1 periportal scores, cluster 2 pericentral scores) and module score values of marker genes from single cell (periportal hepatocytes scores, pericentral hepatocyte scores) using spatial transcriptomics count-data. **b)** Spearman correlations between module score values of markers from spatial transcriptomics data (cluster 1 periportal scores, cluster 2 pericentral scores) and module score values of markers from single cell (periportal hepatocytes scores, pericentral hepatocyte scores) using mouse cell atlas (MCA) data [39].

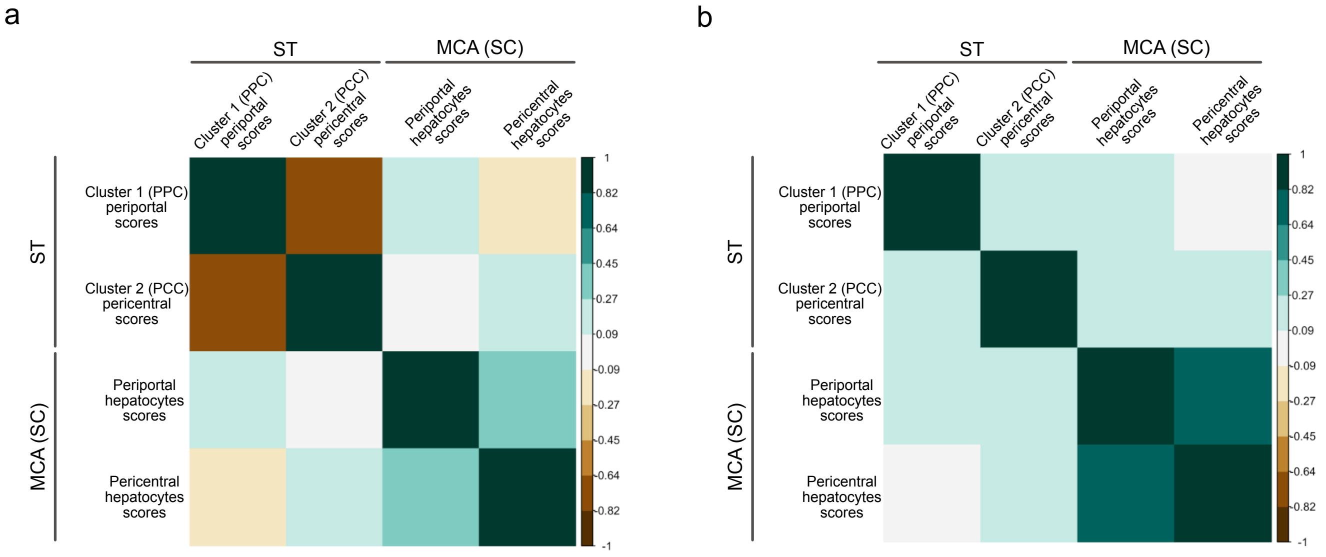


**Supplementary table 3.1:** Spearman correlation values of module score values of periportal and pericentral gene markers from single cell data (hep.pp_MCA, hep.pc_MCA) and module score values of pericentral and periportal markers from spatial transcriptomics data (cluster1_ST, cluster 2_ST) using spatial transcriptomics data.

| | cluster1_ST | hep.pp_MCA | cluster2_ST | hep.pc_MCA |
|---|---|---|---|---|
| **cluster1_ST** | 1 | 0.237944152960821 | -0.747074463978828 | -0.150505061278013 |
| **hep.pp_MCA** | 0.237944152960821 | 1 | -0.0108272714188795 | 0.44290617681838 |
| **cluster2_ST** | -0.747074463978828 | -0.0108272714188795 | 1 | 0.150868310269106 |
| **hep.pc_MCA** | -0.150505061278013 | 0.44290617681838 | 0.150868310269106 | 1 |

**Supplementary table 3.2:** Spearman correlation values of module score values of periportal and pericentral gene markers from single cell data (hep.pp_MCA, hep.pc_MCA) and module score values of pericentral and periportal markers from spatial transcriptomics data (cluster1_ST, cluster 2_ST) using mouse cell atlas (MCA) data.

| | cluster1_ST | hep.pp_MCA | cluster2_ST | hep.pc_MCA |
|---|---|---|---|---|
| **cluster1_ST** | 1 | 0.211384034296339 | 0.0991093609396526 | 0.0515614925278205 |
| **hep.pp_MCA** | 0.211384034296339 | 1 | 0.245429161314913 | 0.674256868173379 |
| **cluster2_ST** | 0.0991093609396526 | 0.245429161314913 | 1 | 0.110786078630394 |
| **hep.pc_MCA** | 0.0515614925278205 | 0.674256868173379 | 0.110786078630394 | 1 |

**Supplementary figure 5:** Pearson correlation of marker genes of all clusters ordered by the first principal component (corrplot package (v.0.84), see methods).

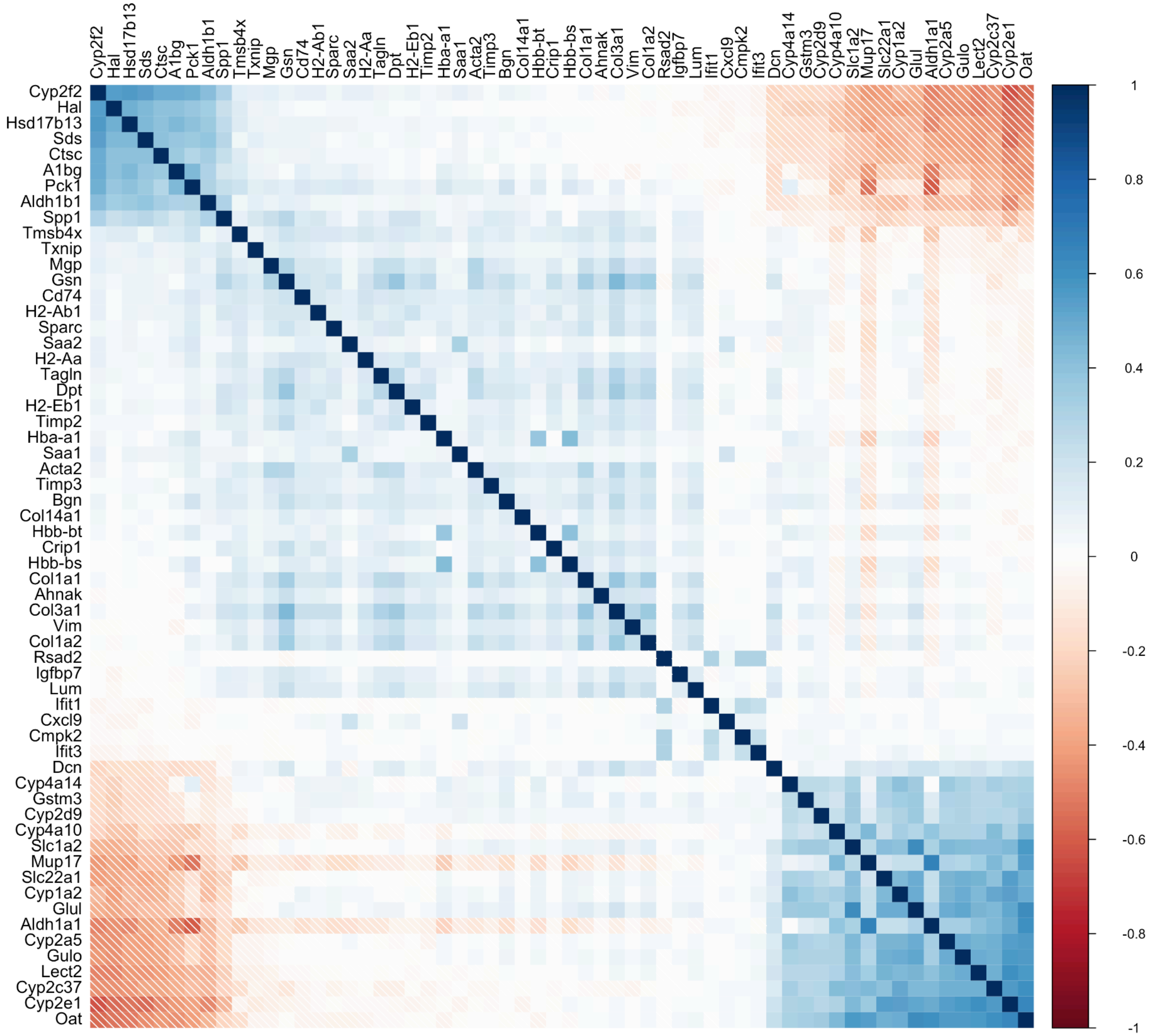

**Supplementary figure 6:** Spatial autocorrelation of marker genes. Bar plot of genes with spatial autocorrelation above a threshold of 0.2 for spatial autocorrelation. Higher values indicate stronger spatial correlation of the gene, while lower values indicate a more random distribution of the gene across the tissue. The color refers to the identified cluster identity.

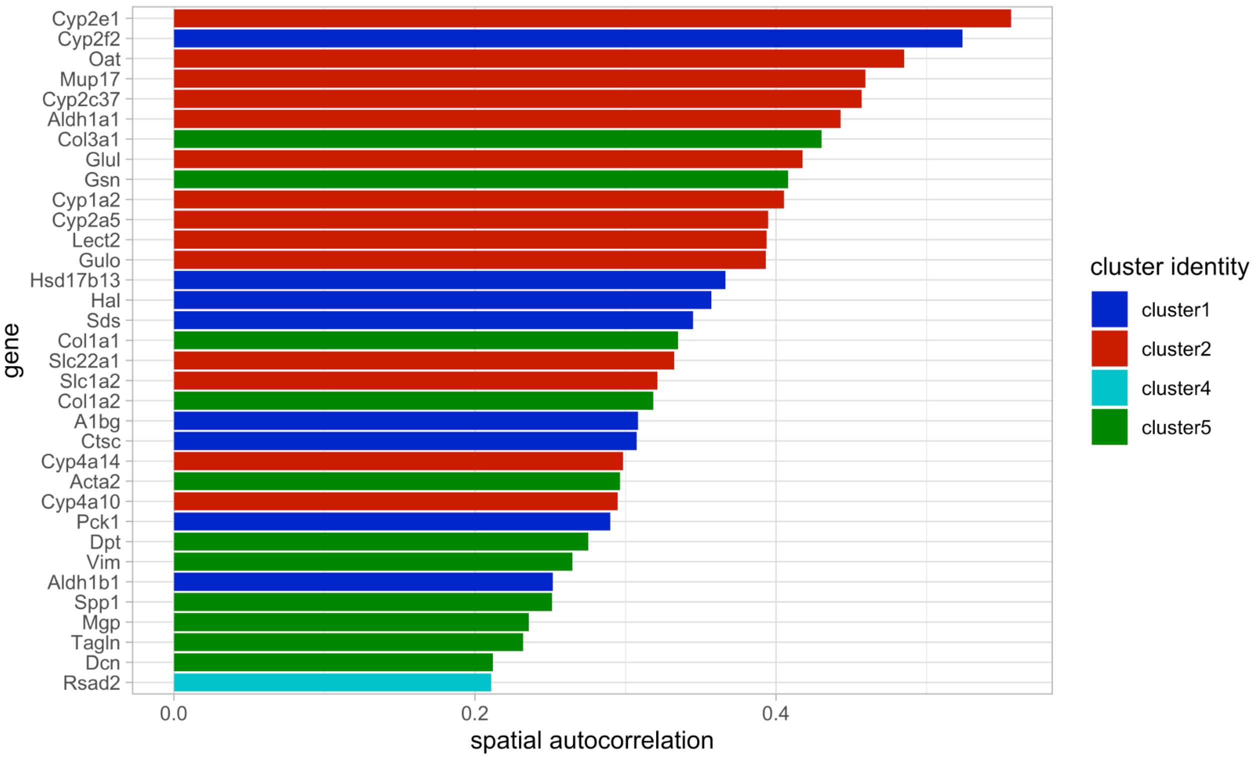

**Supplementary figure 7.1:** Visualization of marker genes of periportal annotated cluster 1. **a** *Sds* expression **b** *Cyp2f2* expression,and **c** module scores (see methods) for periportal markers of cluster 1 (bottom) on Hematoxylin- and Eosin-stained tissue for all sections of replicate 1 to replicate 3. Descending expression values/scores is visualized by decreasing opacity of spots.

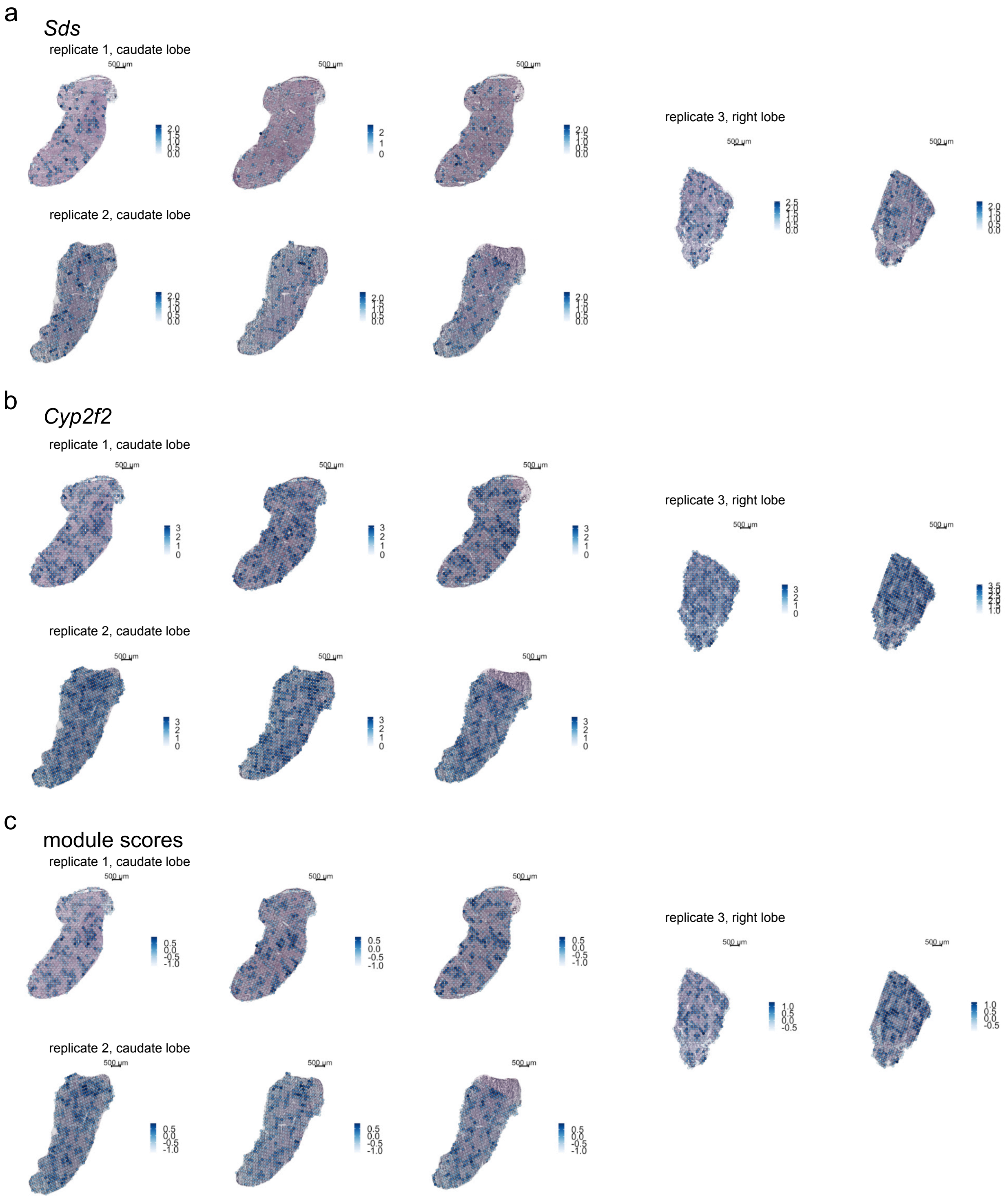

**Supplementary figure 7.2:** Visualization of marker genes of pericentral annotated cluster 2. **a** *Glul* expression **b** *Cyp2e1* expression,and **c** module scores (see methods) for pericentral markers of cluster 2 (bottom) on Hematoxylin- and Eosin-stained tissue for all sections of replicate 1 to replicate 3. Descending expression values/scores is visualized by decreasing opacity of spots.

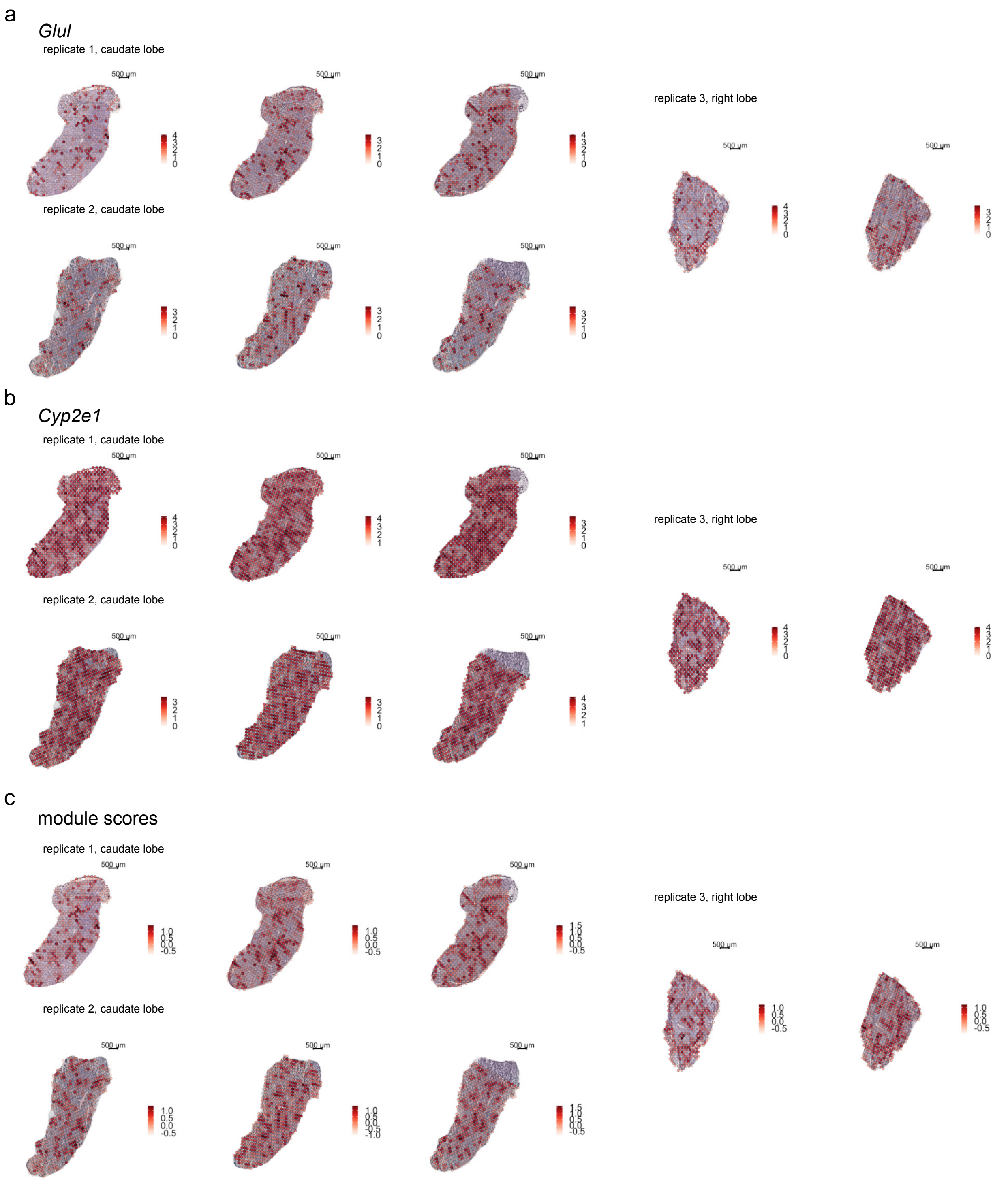

**Supplementary figure 8:** Visualization of expression of **a** periportal and **b** pericentral marker genes identified by unsupervised clustering and DGEA of ST data across reconstructed spatial layers (1-9) of single cell data zonation matrix (Halpern et al., 2017). In combined plots, single cell expression data was scaled to the maximal expression value in the group of genes (see methods). Each periportal and pericentral marker gene is additionally plotted individually with original expression values of single cell data.

a

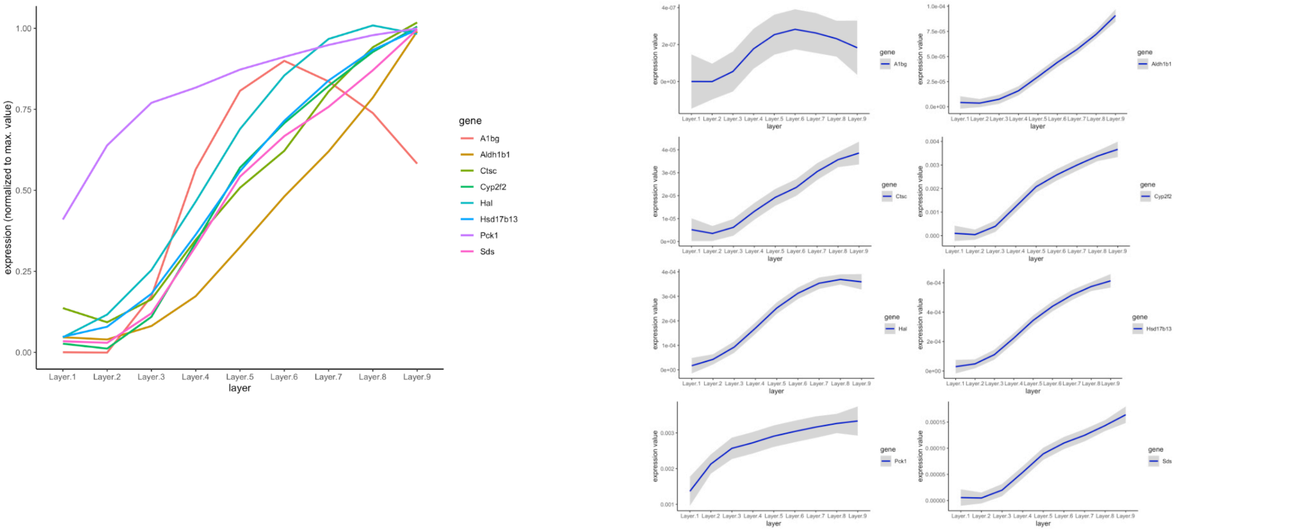


b

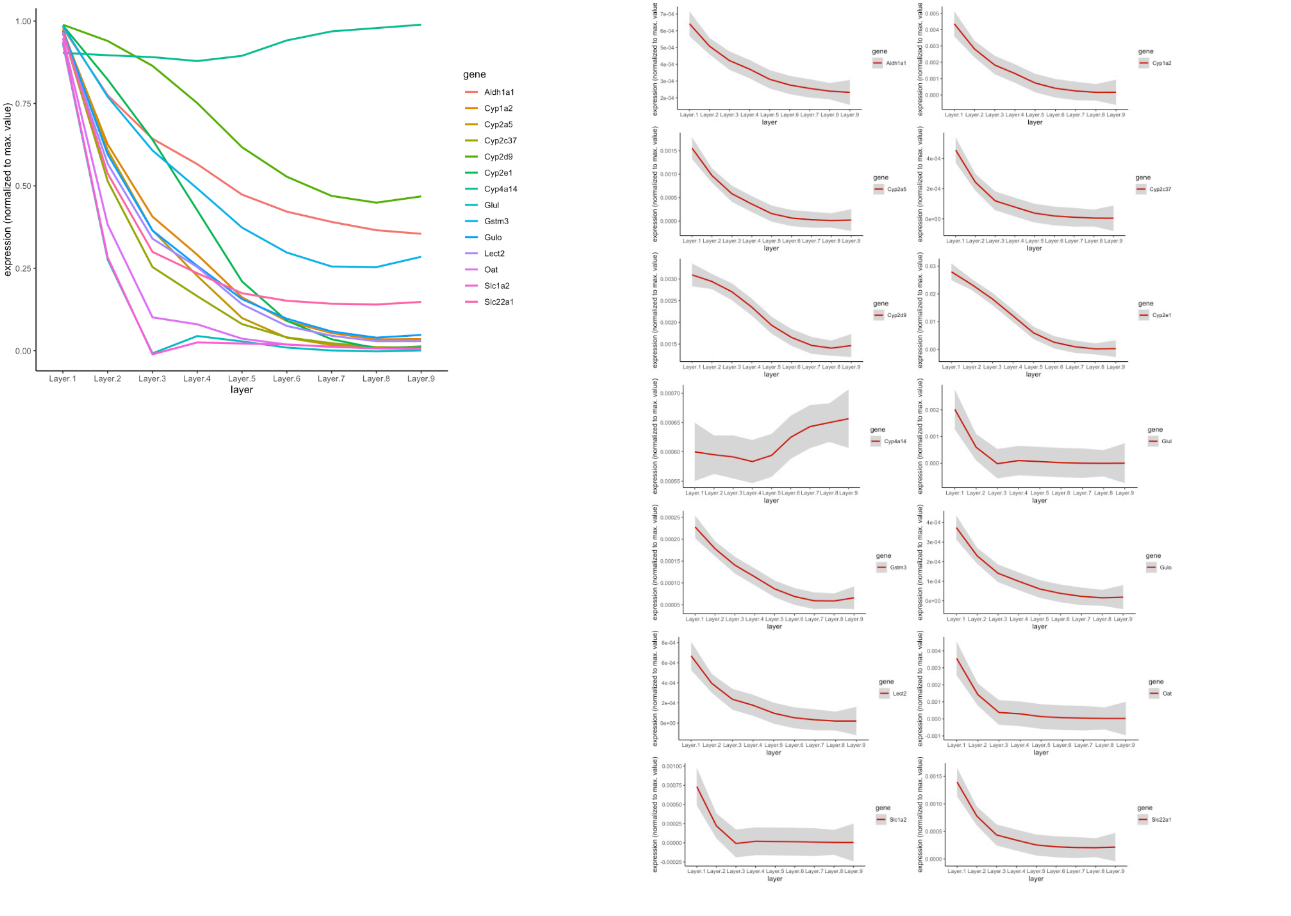

**Supplementary figure 9:** Expression by distance of annotated cell types of MCA single cell data from outer portal and central vein borders. Distances of pericentral hepatocytes from central (C) to portal (P) veins are depicted in the figure (on the left). Distances of periportal hepatocytes from portal (P)to central veins (C) are depicted in the figure (on the right).

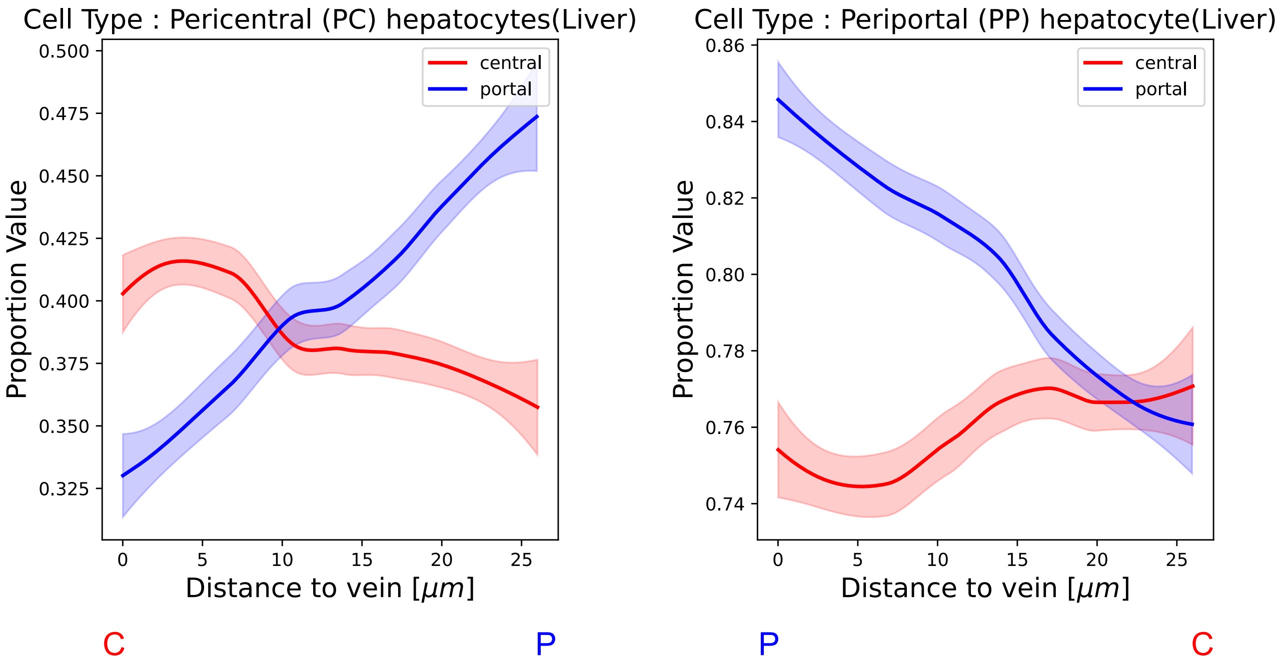

**Supplementary figure 10:** Visualization of spots under the tissue, assigned to cluster 5 by unsupervised clustering and visualization on all HE stained tissue sections. **a)** Replicate 1 with 3 sections of a part of a caudate lobe. **b)** Replicate 2 with 3 sections of a part of a caudate lobe and **c)** replicate 3 with 2 sections of a part of a right lobe.

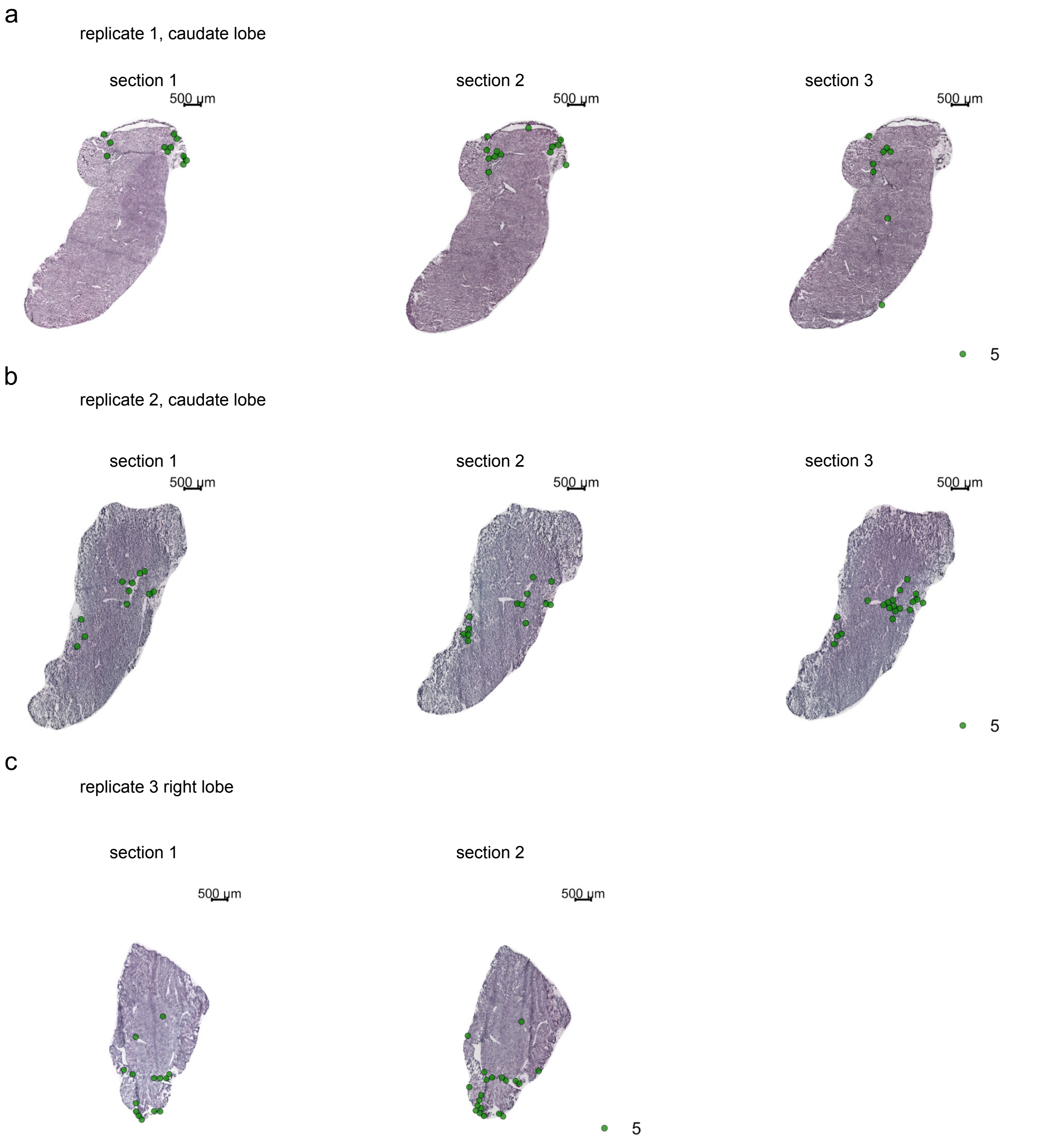

**Supplementary figure 11:** Visualization of four marker genes annotated to cluster 5 showing the highest positive logFC within the cluster: **a** *Vim,* **b** *Col3a1,* **c** *Col1a2* and **d** *Gsn* superimposed on all HE stained tissue sections of replicate 1 to replicate 3. Descending expression values/scores are visualized by decreasing opacity and a change from lighter to darker color.

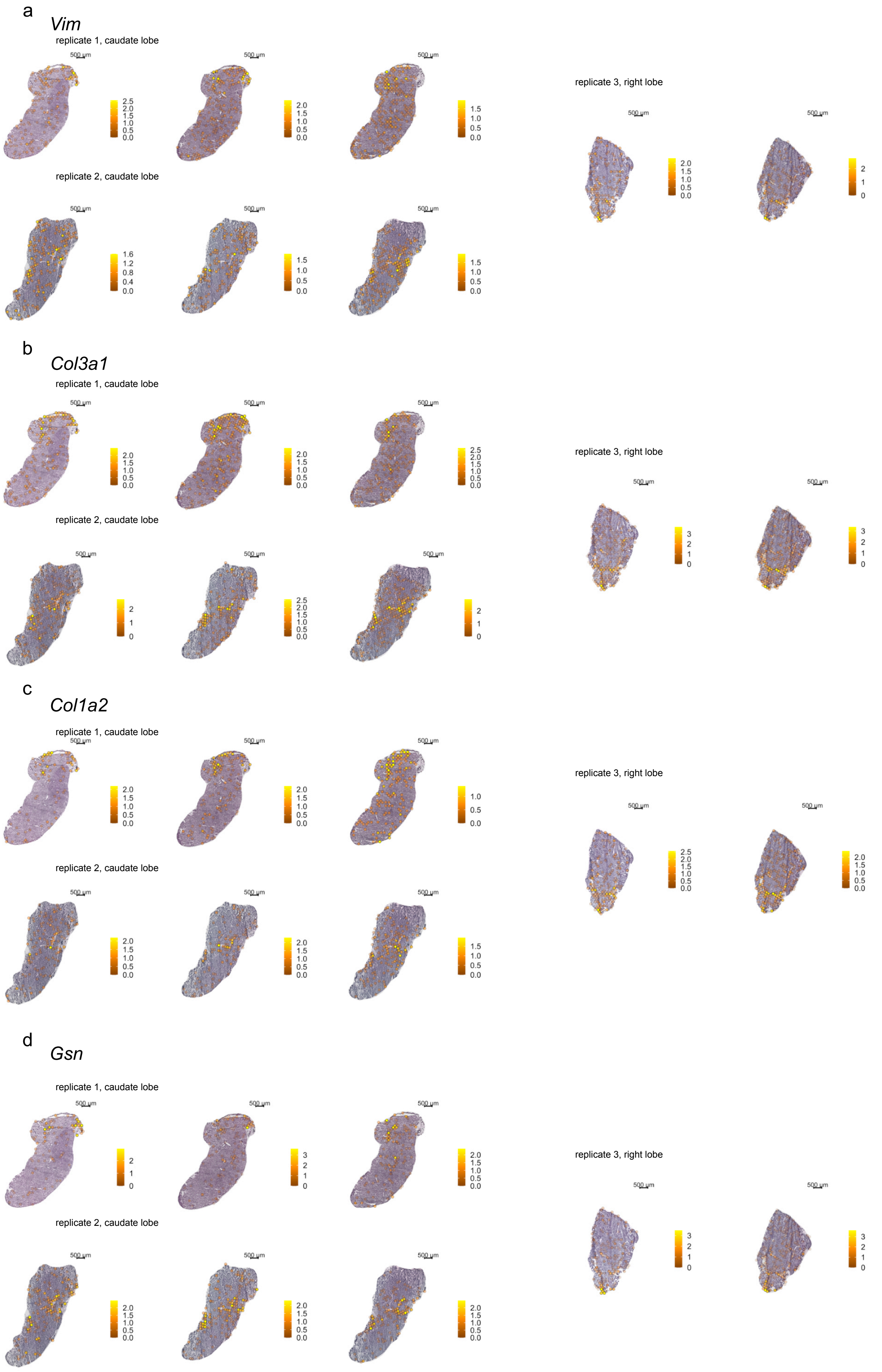

**Supplementary figure 12:** Visualization of pathway analysis on tissue sections.**a** Module scores for gene-sets with GO:0030199: “collagen fibril organization” annotation and **b** GO:0034097: ”response to cytokine” (bottom), superimposed on all tissue sections and replicates. Descending expression values/scores are in addition to change color visualized by decreasing opacity of spots.

a

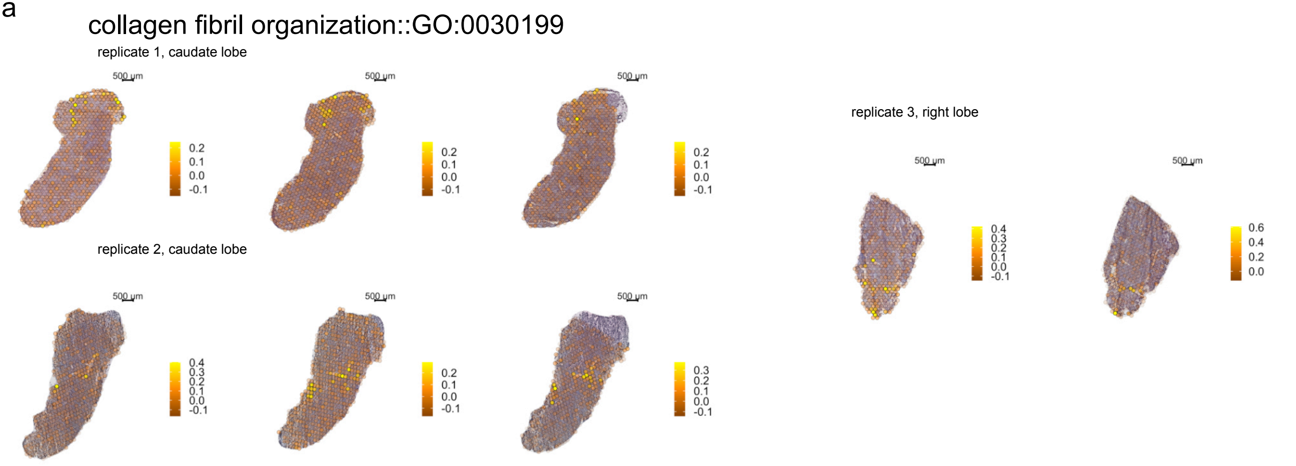


b

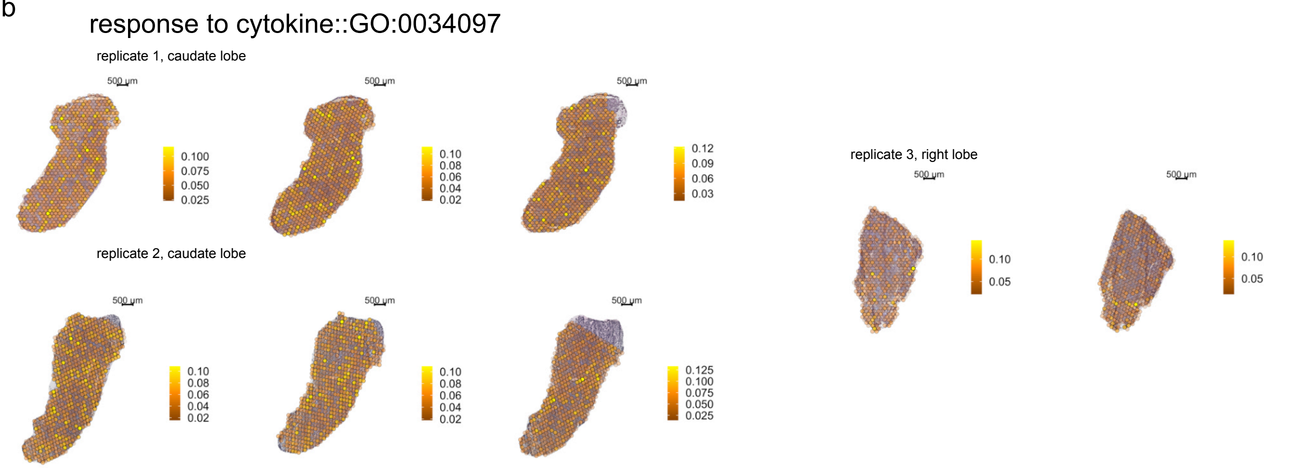

**Supplementary figure 13:** Shortlist of five marker genes of cluster 1 (portal vein) and cluster 2 (central vein) identified by unsupervised clustering of ST data. The central and portal markers depicted here, exhibit the highest logFC for the respective cluster in the ST data and were used in computational analyses of expression by distance plots and the computational annotation of vein types in the tissue.

| central markers | portal markers |
|---|---|
| Glul | Sds |
| Oat | Cyp2f2 |
| Slc1a2 | Hal |
| Cyp2e1 | Hsd17b13 |
| Cyp2a5 | Aldh1b1 |
